# Charting Single Cell Lineage Dynamics and Mutation Networks via Homing CRISPR

**DOI:** 10.1101/2024.01.05.574236

**Authors:** Lin Wang, Wenjuan Dong, Zheng Yin, Jianting Sheng, Chika F. Ezeana, Li Yang, Xiaohui Yu, Solomon SY Wong, Zhihao Wan, Rebecca L. Danforth, Kun Han, Dingcheng Gao, Stephen T. C. Wong

## Abstract

Single cell lineage tracing, essential for unraveling cellular dynamics in disease evolution is critical for developing targeted therapies. CRISPR-Cas9, known for inducing permanent and cumulative mutations, is a cornerstone in lineage tracing. The novel homing guide RNA (hgRNA) technology enhances this by enabling dynamic retargeting and facilitating ongoing genetic modifications. Charting these mutations, especially through successive hgRNA edits, poses a significant challenge. Our solution, LINEMAP, is a computational framework designed to trace and map these mutations with precision. LINEMAP meticulously discerns mutation alleles at single-cell resolution and maps their complex interrelationships through a mutation evolution network. By utilizing a Markov Process model, we can predict mutation transition probabilities, revealing potential mutational routes and pathways. Our reconstruction algorithm, anchored in the Markov model’s attributes, reconstructs cellular lineage pathways, shedding light on the cell’s evolutionary journey to the minutiae of single-cell division. Our findings reveal an intricate network of mutation evolution paired with a predictive Markov model, advancing our capability to reconstruct single-cell lineage via hgRNA. This has substantial implications for advancing our understanding of biological mechanisms and propelling medical research forward.

## Introduction

Lineage tracing is a powerful technique for mapping the dynamic process of cell division, and various strategies to reconstruct cell lineage tracing have been developed^1–10^. CRISPR-Cas9 genome editing has been integrated into lineage tracing techniques, leveraging its ability to introduce diversity and irreversible mutations at specific genomic loci that can be inherited by subsequent generations, thereby enabling the differentiation of cells within distinct lineage branches^11–15^. Numerous studies in developmental progeny tracing have utilized these techniques^16–19^. Recently, these studies have advanced to the single-cell resolution, combining with single-cell transcriptomics to enhance phylogenetic trees through the inclusion of information acquired through single-cell tanscriptomics^20–24^.

Several studies have also applied these techniques to map the clonal evolution of cancer metastasis^25–27^. *Quinn et al.* created a Cas9-based lineage tracer and employed single-cell sequencing to trace phylogenies and track the movement of metastatic human cancer cells implanted in the lung of a mouse xenograft model. In this setting, they found that cancer cells exhibited diverse metastatic phenotypes and subclones exhibited differential gene expression profiles, some of which were previously associated with metastasis^25^. *Simeonov et al.* developed macsGESTALT, a CRISPR-Cas9-based lineage recorder simultaneous capturing both lineage and transcriptomes for single cells, and lineage reconstruction in a metastatic pancreatic cancer model reveals extensive bottlenecking and subpopulation signaling, as well as specific transcriptional states associated with metastatic aggression and predictive of worse outcomes in human cancer^26^.

The CRISPR-Cas9 system includes a single-guide RNA (sgRNA) that guides the Cas9 protein to recognize and cleave a specific genomic locus, resulting in a Cas9-induced double-stranded DNA break^28^. Subsequently, mutations are introduced at the targeted locus through the non-homologous end joining (NHEJ) repair process^29^. *Shen et al.* suggests that the possibility to predict the outcomes of template-free end-joining repair, a common DNA repair mechanism used by cells to fix double-stand breaks induced by CRISPR-Cas9^30^. While sgRNA-based CRISPR-Cas9 system has proven versatile and widely applicable, a modified version of the CRISPR-Cas9 system, homing CRISPR, recently has been proposed as an alternative over the typical sgRNA system^31, 32^. A homing CRISPR system employs a homing guide RNA (hgRNA) to direct Cas9–gRNA nuclease activity to the guide RNA locus itself, simplifying the system by linking the guide RNA locus and the target site into one entity.

To engineer hgRNA, a specific modification is made to the gRNA spacer. In this case, the immediate downstream of *S. pyogenes* gRNA spacer mutates from ‘GUU’ to ‘GGG’ (Fig. 1a). This spacer modification endows the gRNA with the capability for multiple rounds of editing, i.e., the system allows for repeated targeting and editing of the same genomic locus. This enables retargeting and evolvability of barcodes, offering advantages including generation of more diversity in edited sequences compared to the established CRISPR-Cas9 system, lower susceptibility to large deletions that could erase barcode or genetic information, imposing a lower burden on cells, and minimizing disturbances to developmental processes^33,34^. Studies have utilized the hgRNA system to investigate bone metastasis in breast cancer; yet despite the advantages of hgRNA, it has not reached single cell lineage tracing coupled with single cell transcriptomics^35,36^.

**Fig. 1.**
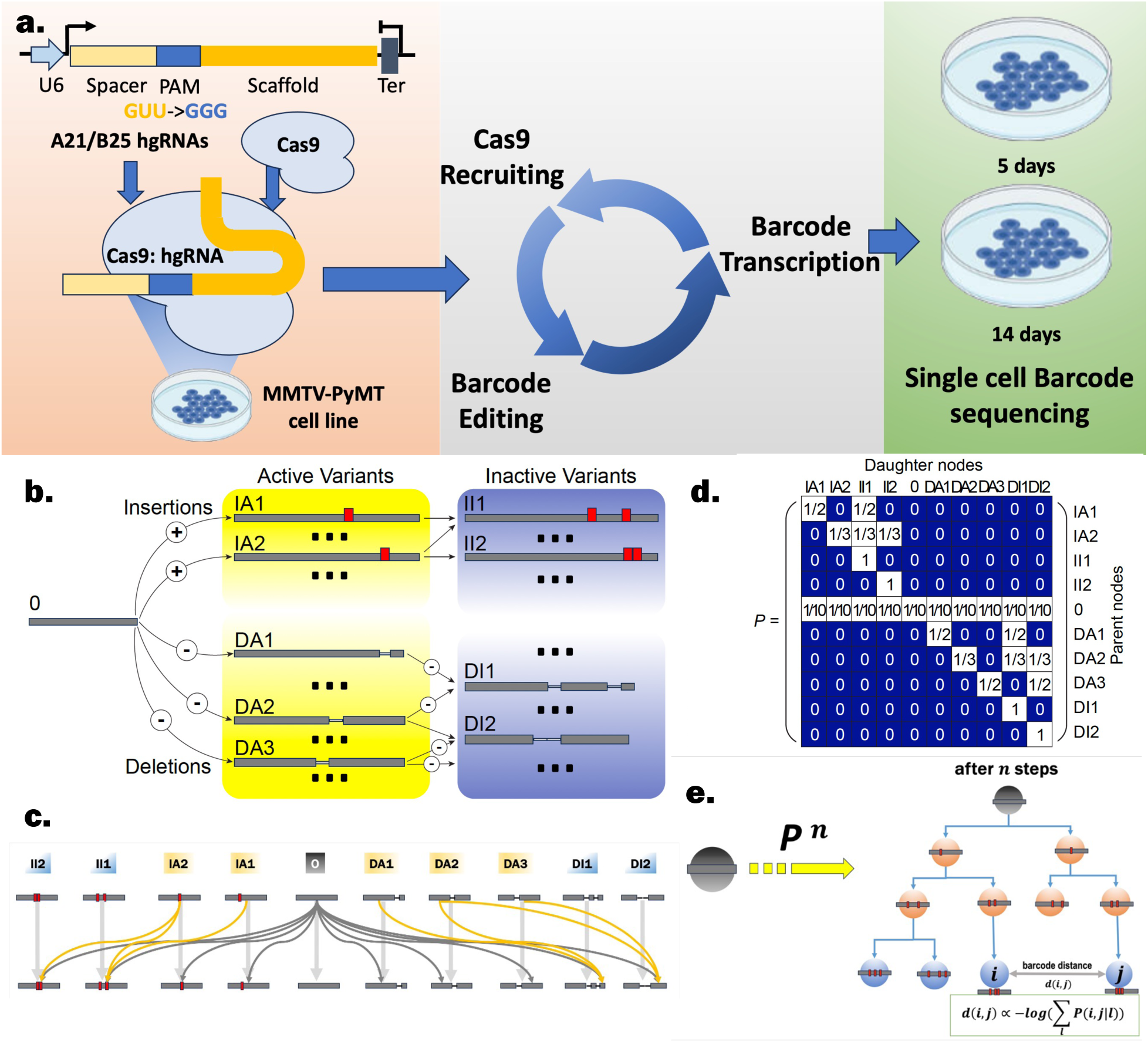
LINEMAP characterizes the specific paths of hgRNA mutations, elucidating single-cell lineage relationships. (a) MMTV-PyMT cell line with A21 and B25 hgRNA barcodes was cultured for 14 days. The specifically designed PAM section (GUU->GGG) in the hgRNAs ensures continuous cycles of Cas9 complex recruiting, barcode editing and barcode transcriptions, chronicled with cell cycles. Single cell barcode sequencing from cells on day 5 and day 14 recorded the resulted barcode lineage. (b) Illustration of the mutation evolution network: Each node represents a specific mutation allele. Every mutation’s downstream mutations must contain the exact mutation at every base. Active variants can undergo further mutations downstream, while inactive variants remain unchanged in all offspring. (c) The deduced transition diagram for the Markov model: Mutations can transition to one another if they are connected by a directed edge. (d) The learned stochastic matrix for the Markov model: The probability *P*_*i,j*_ should be zero if there is no way for mutation *i* to transition to mutation *j*. (e) Calculation of the distance between two cells based on their barcodes using the stochastic matrix of the Markov model: Here, *i* and *j* represent two barcodes from distinct cells, and *P*(*i*, *j*|*l*) represents the probability of the two daughters’ barcodes being *i* and *j* respectively, given that their parent cell’s barcode is *l*.

A featured characteristic of hgRNA self-retargeting allows for multiple runs of a specific mutation at the genomic locus. The process continues until the mutation disrupts the original sequence’s structure, leading to the formation of a stable inheritable mutation that can be passed on all future generations. Nonetheless, the intermediate evolutionary pathways during these multiple cycles of evolution are intricate and nebulous. There is a possibility that distinct evolutionary paths could eventually mutate into the same mutation alleles, thereby increasing the difficulty to identify the previous parent mutation allele. This complexity obfuscates the ability to accurately trace the parent mutation allele or reconstruct the exact sequence of events that led to a particular mutation. Furthermore, unlike sgRNA, which produces irreversible and cumulative mutations, the hgRNA system may not exhibit the equivalent level of irreversibility. Uncertainty regarding the irreversibility of hgRNA mutations can impact the predictability and stability of the system over multiple editing cycles. These limitations associated with hgRNA may hinder its potential for tracing the development history and lineage of individual cells within more complex biological systems or organisms.

In this study, we present LINEMAP (LINeage of Evolving Mutations and Phenotypes), a computational modeling approach to precisely characterize the path-specific nature of hgRNA mutations. LINEMAP introduces the concept of the Mutation Evolution Network, a graphical model to illustrate the relationships among different genetic mutations or variants over time (Fig. 1b). We hypothesize that mutation alleles can only evolve along the paths in the network without reversing, and these mutation alleles become more complex as additional mutations accumulate over time. We then employ a Markov process model to elucidate the evolution pathways of hgRNA (Fig. 1c). Whether two mutation alleles can transition is determined by the mutation evolution network. Next, we compute a maximum-likelihood estimate of the transition probability within the stochastic matrix for the Markov model (Fig 1d). Finally, we develop a novel reconstruction algorithm using statistical method based on the Markov model to estimate the lineage relationships in single cells (Fig. 1e).

We developed an A21B25 barcode MMTV-PyMT (mouse mammary tumor virus-polyoma middle tumor-antigen)-Cas9 tracing cell line (PyMT-Cas9 tracing cell line), enabling us to investigate the mutation evolution network’s ability to characterize the mutation evolution of hgRNA alleles within cell lines (Fig. 1a). We conducted both computational (in silico) and laboratory-based (in vitro) experiments to evaluate our hypothesis of irreversibility and cumulativeness in hgRNA mutations. The results from all experiments consistently demonstrated our capability to reveal the actual mutation evolution pathways of hgRNA, further confirming the irreversible and cumulative nature of hgRNA mutation evolution. Leveraging these characteristics, we can employ hgRNAs to reconstruct lineage tracing work, such as progenitor mapping, at the single-cell level.

## Results

### An analytic pipeline to identify the mutation alleles in the PyMT-Cas9 tracing cell line

A21 and B25 are two homing guide RNA (hgRNA) barcodes utilized in our PyMT-Cas9 tracing cell line. Each of these barcodes contains a unique spacer with lengths of 21 and 25, respectively^35^. The barcode editing for the PyMT-Cas9 tracing cell line was initiated by adding doxycycline at the concentration of 500 nM/ml (Methods). We harvested the cells at 5 days and 14 days after initiation of hgRNA editing (Fig. 1a). For this study, we developed a sequencing data processing pipeline for the PyMT-Cas9 tracing cell line data to facilitate the identification of mutation alleles for each hgRNA barcode (Fig. 2a and method). In total, we collected 2,677 cells on day 5 after treatment of doxycycline, resulting in 169 unique mutation alleles observed for A21, and 3,564 cells on day 14 post doxycycline induction, resulting in 257 unique mutation alleles observed for A21. Combining data from these two time points, we identified a total of 331 unique mutation alleles, including the original hgRNA barcode (Fig. 2b and Supplementary Table 1). Mutation frequencies were calculated based on the number of cells in which they were observed. Of these 331 mutations, insertion-type mutations were predominant at 61.6%, but were only observed in 16.2% of all cells. Conversely, deletion-type mutations had higher frequencies, with 83.5% of cells displaying such mutations. Among deletion-type mutations, position −6 is most likely to be deleted, and the likelihood of being deleted decreases sequentially on both sides of −6 (Fig. 2c). In insertion-type mutations, the most common insertion position is located three bases upstream of the PAM sequence, accounting for 46.7% of all insertion-type mutations (Fig. 2d). These results corresponded to the location of double-stranded DNA breakage and thus well within expectations. Furthermore, at this position, insertions of 1 or 2 bps were observed at a high frequency, while insertions at other positions tend to have longer sequence lengths (Fig. 2e and Supplementary Fig. 1).

**Fig. 2.**
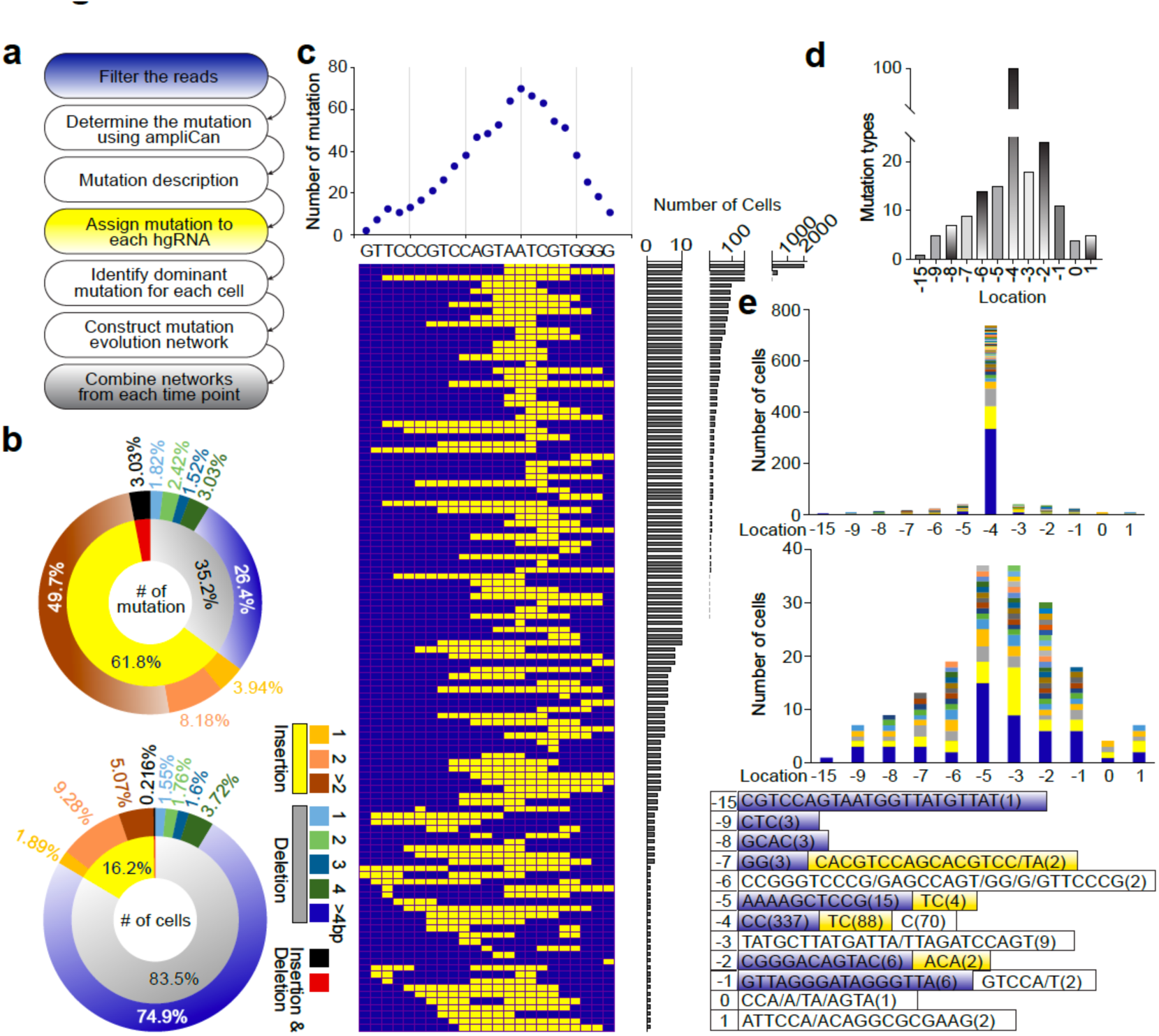
Overview of mutation alleles for hgRNA A21 identified in the PyMT-Cas9 tracing cell line. (a) Workflow of the analytic pipeline used to identify the mutation alleles. After filtering and alignment steps, each candidate mutation allele will be stored in a descriptive matrix that records the mutation’s location, type (insertion and/or deletion), and the exact differences of each base compared to initial hgRNA barcode. This storage structure will benefit our subsequence analysis, such as constructing the mutation evolution network. (b) Top: the percentage of each category of mutation alleles in the total mutation count for hgRNA A21 in the PyMT-Cas9 tracing cell line. Bottom: the percentage of each category of mutation alleles in the total number of cells for hgRNA A21 in the PyMT-Cas9 tracing cell line. (c) All the deletion-type mutation alleles for hgRNA A21 identified in the cells. Each row corresponds to a unique mutation sequence with yellow spots indicating deletions, blue spots indicating intact bases. The frequency at which each base is deleted in plotted at the top, and the sequence includes the hgRNA A21 and the 3-bp PAM. The total number of cells in which these are observed is indicated on the right. (d) The frequency of the insertion positions among all insertion mutations. (e) The frequency of insertion lengths at each position.

### Construct mutation evolution network

We disregarded any mutation alleles observed in fewer than 10 cells at both time points in further analysis due to the low observation frequency, introducing statistical noise. We were left with 54 different mutation alleles (Supplementary Table 1). We use these mutations to construct a mutation evolution network according to the following principles. First, the original intact barcode sequence serves as the root node. Second, for each node, its immediate daughter node corresponds to a mutation allele that retains all the same insertions and deletions (indels) at the same location while possessing additional indels at other bases in the sequence. Third, no other mutation allele can function as both the parent node’s daughter and the daughter node’s parent (Fig. 1b).

These guidelines allows us to conceptualize the mutation evolution in order of complexity. It is certain that each parent node could generate multiple daughter nodes, and conversely, each daughter node might have more than one parent node, suggesting that multiple paths lead to the same mutation alleles. We developed a methodology to distinguish between active mutation alleles that can be retargeted and continuously edited from inactive alleles that cannot be further altered. By comparing the frequency of each mutation allele at two time points, we can determine their status. If the frequency increases, it indicates an inactivated allele, suggesting that more cells have transitioned to that state. Conversely, the decreased frequency suggests an active allele, implying that more cells have transitioned from a parent mutation to other mutation alleles. The mutation alleles identified as active all have a left sequence no shorter than 16 base pairs (bps), which is consistent with experimental findings that Cas endonuclease will not show any activity when gRNA sequences are shorter than 16 bps^37,38^.

### Modelling the mutation evolution using Markov Process model

Next, we model the mutation evolution path using a Markov process. We define each mutation allele as a state in the Markov model, one cell division corresponding to one transition in states. Mutations occur during cell division or unchanged with each cell division. Based on the mutation evolution network, we can define the transition diagram for the Markov model as follows: for each active mutation, there is a possibility of transitioning to any mutation allele downstream in the mutation evolution network, the equivalent to a transient state in the Markov model. Taking the root node as an example, the root node has the potential to transition to any mutation allele. For the inactive mutation allele, it transitions only to itself, which represents an absorbing state in the Markov model (Fig. 1c). Thus, the transition probability in the stochastic matrix of the Markov model has many values of 0 (indicating no transition can occur between those two mutation alleles) and 1 (representing an inactive mutation allele that transitions only to itself) (Fig. 1d). We then computed a maximum likelihood estimate of the transition probability in the stochastic matrix for values that are not 0 or 1. This process involves solving a nonlinear optimization problem (Methods).

### The methodology inferred the mutation evolution with high accuracy in silico

Transition probabilities represent the likelihood of a mutation allele changing or transitioning to another allele during a specific event or process. Various factors can influence these probabilities, such as the mutational mechanisms, selection pressures, or random genetic drift^39^. To verify the accuracy of the methods outlined above in revealing actual mutation transitions and estimating transition probabilities, we synthesized three sets of mutation evolution networks, each with different numbers of mutations, and used this network to generate mutation alleles for 10000 cells. The number of mutations in each network gradually increases: 20, 30, and 100 mutations, including the unmutated hgRNA, respectively (Fig. 3a, 3b). Active mutation alleles are defined as the closest 20% of nodes to the root node in each network. For each active mutation allele, the transition probability to all the mutation alleles it could transition to is randomly assigned within the stochastic matrix.

**Fig. 3.**
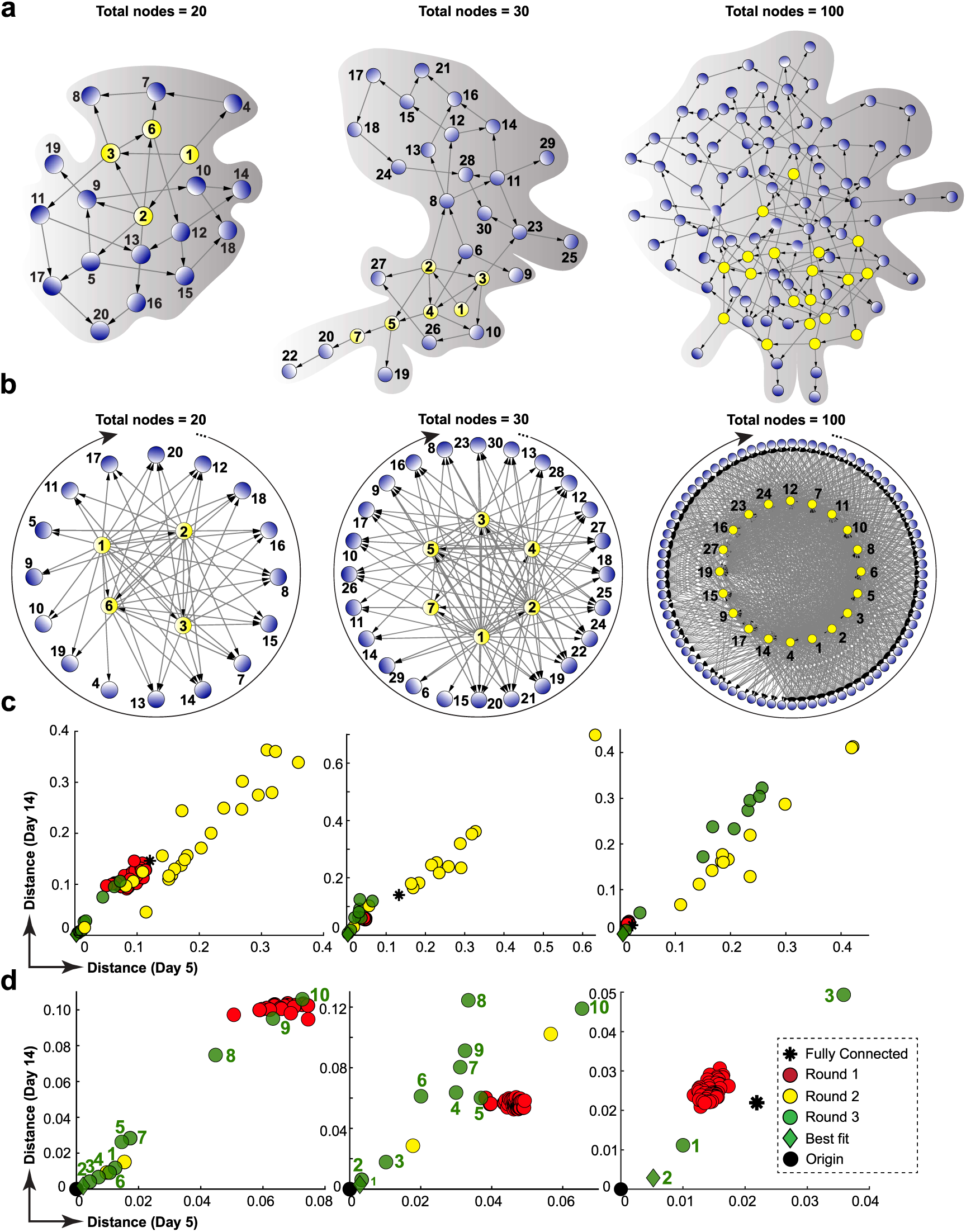
In silico experiments to validate our methodology to infer the accurate mutation evolution. (a) Synthetic mutation evolution network with 20, 30, and 100 mutation alleles. The node in yellow indicates an active mutation, and node in blue indicates an inactive mutation. (b) Transition diagrams deduced from the generated mutation evolution networks. The stochastic matrix of the Markov model determines the mutation alleles for 10,000 simulated cells. (c) Distance measurements between frequency vectors resulting from three experimental rounds and the actual frequency vector for simulated data. The X-axis represents the distance at day 5, and the Y-axis at day 14. Red, yellow, and green points denote the results of experiments 1, 2, and 3. (d) Zoomed-in section of figure c displaying the results of experiment 3. Numbers 1-10 represent various proportions of specified active mutations from 10% to 100%. The number 2 indicates a perfect match between specified active mutations and those used in the actual simulated data, at 20%.

To ensure the probabilities are sufficiently high for effective model learning, a minimum probability threshold of greater than 0.01 is imposed. We simulate cell divisions based on the cell cycle of the PyMT-Cas9 tracing cell line (method). We randomly simulate the number of cell divisions undergone by 10,000 cells at specific time points (5-day and 14-day).

Subsequently, we calculate the proportion of cells with varying numbers of divisions. These 10,000 cells progress through Markov state transitions based on their division history. The final state, determined after the Markov process, represents the mutation allele of each cell. Finally, we tally the number of cells for each mutation allele and calculate a frequency vector for all mutation alleles within the population at each time point. Our method thus combines computational modeling, simulation, and probabilistic modeling to explore evolution networks and transition probabilities of hgRNA mutations in a controlled setting.

The observed mutation evolution network implies a premise of irreversibility and seems to align with the experimental observations^35^. All active mutations, while still capable of undergoing further mutations, can only involve adding additional insertions and deletions elsewhere based on the existing indels. It is not possible for the original mutations to be eliminated, i.e., to revert to the mutation state of their parent node or even any upstream node.

To test this hypothesis, we conducted three rounds of experiments. The first round aimed to verify the assumption of irreversibility. For each group with a different number of mutation alleles, we randomly determined whether one mutation allele can transition to another. This set up may lead to the emergence of cyclic structures in the network or cases where two mutation alleles can mutually transition. We accounted for the extreme scenario should all mutation alleles transition into one another, which creates a fully connected network in the transition diagram.

In the second round of experiments, we adhered to the assumption of irreversibility while focusing on validating our mutation evolution network-based transition diagram. We randomly generated several mutation evolution networks with the specified number of mutation alleles, ensuring that the root node represented the original barcode and formed a directed acyclic graph. Active mutations were defined as the closest 20% of nodes to the root node. We deduced transition diagrams from these generated networks.

In the third round of experiments, we used the mutation evolution network that the simulated 10,000 samples were generated from, but with varying numbers of active mutation alleles. We defined active mutation alleles as nodes closest to the root node within the range of 10% to 100% and generated transition diagrams accordingly.

After generating three rounds of mutation evolution networks and transition diagrams, we compute a maximum likelihood estimate for the stochastic matrix and calculate the frequencies of each mutation allele based on the estimated stochastic matrix and the number of cell divisions. The resulting frequency vector is compared to the real frequency vector for the 10,000 simulated samples generated using the mutation evolution network we synthesized.

This comparison involves calculating the Euclidean distance between the two vectors. In these experiments, our objective is to validate the capability of our method in accurately inferring the correct mutation network and transition diagram, which in turn enables us to correctly deduce the patterns of mutation evolution.

We can achieve mutation allele frequency vectors that closely match the ones obtained from the simulated samples only when the mutation evolution network selected is in complete alignment with the network used for simulating samples. This consistency holds across all three types of networks and emphasizes the irreversibility of the mutation evolution network. Any deviations from our defined network or discrepancies in determining the active alleles result in disparities in mutation frequencies at our chosen time points. To obtain a frequency vector that closely resembles our generated data (Fig. 3c, 3d), it is imperative to first establish the accurate network, identify the correct active mutation alleles, deduce the proper transition diagram, and learn the maximum likelihood transition probabilities. Consistent results are observed when selecting varying proportions of active mutation alleles in the network, ranging from 10% to 40% (Supplementary Fig. 2, 3, 4).

### Characteristic of the mutation evolution of hgRNA A21 in PyMT-Cas9 tracing cell line

Using the methodology mentioned above, we constructed a mutation evolution network for the 54 selected mutation alleles for A21 barcode in our PyMT-Cas9 cell line (Fig. 4a). Among the 54 mutation alleles identified, 13 of them are considered active mutations, determined by comparing the frequencies between two time points (Supplementary Table 1). Across the active mutation alleles, their frequencies decreased from day 5 to day 14, which is in line with the evolution of hgRNA editing barcode. Some inactive mutations had exceedingly low frequencies, often less than 0.005%, thus were challenging to distinguish. The evolution of mutations is visually represented by the evolution network, where the same mutation allele might undergo different paths. This suggests the possibility of diverse lineage origins. In addition, we proceeded to compare the cell counts associated with each mutation allele and their frequencies at the two time points (Fig. 4a). There is evidence suggesting that mutation alleles along certain pathways have a higher number of cells, indicating a higher probability of mutations evolving in that direction. Furthermore, mutations with lower complexity scores, or active mutation alleles, exhibit higher frequencies at day 5, while mutations with higher complexity scores, or inactive mutation alleles, display higher frequencies at day 14. Complexity score is simply defined as adding 1 for each base deletion or insertion. This consistent trend of accumulation is observed throughout the entire mutation evolution network.

**Fig. 4:**
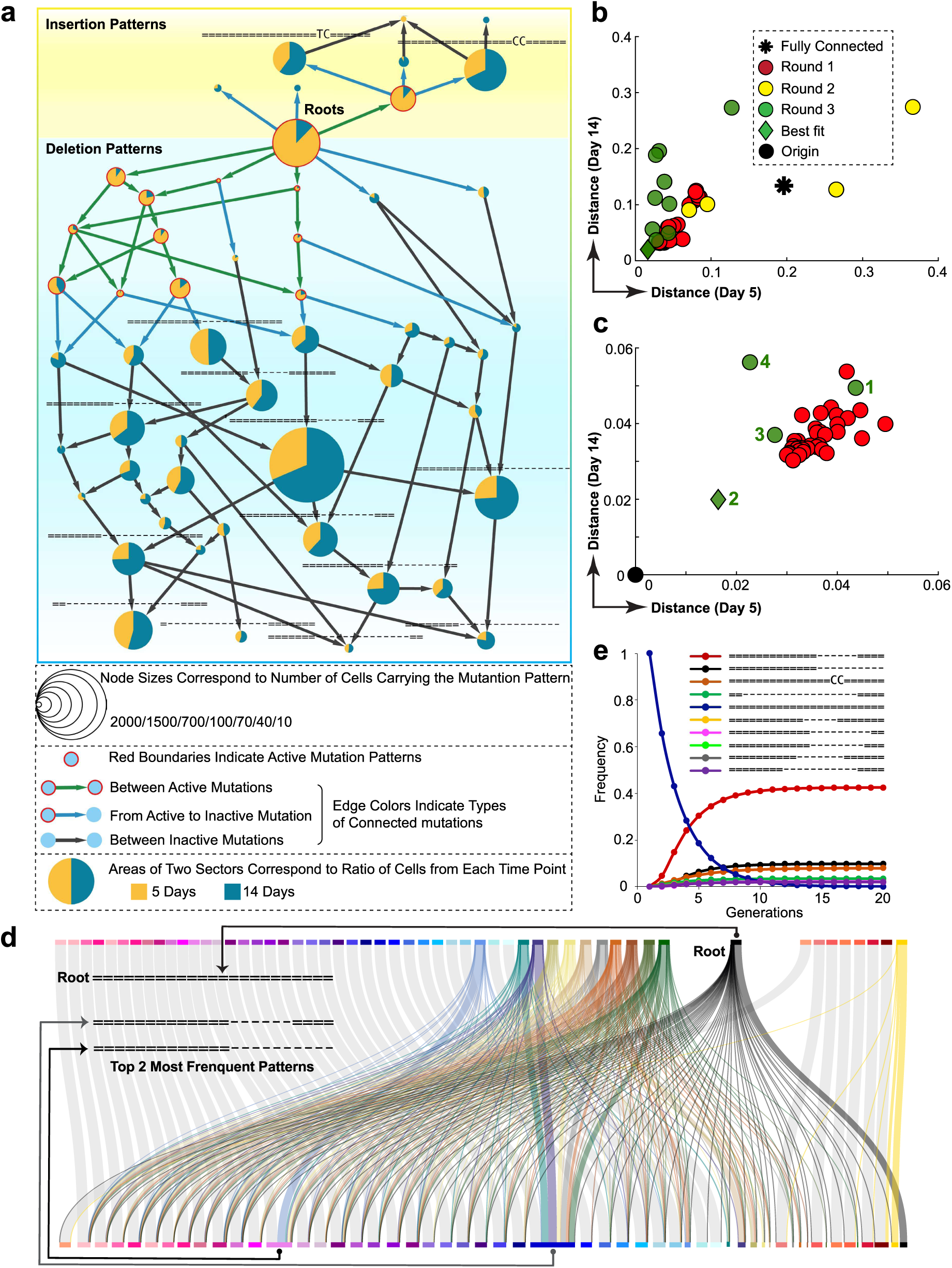
Validation of the methodology against the actual frequency vectors obtained in the PyMT-Cas9 tracing cell line. (a) The mutation evolution network for the 54 mutation alleles of hgRNA A21 in the PyMT-Cas9 tracing cell line. The mutation evolution network integrated with the cell numbers at each time point. The size of each node represents the number of cells observed across two time points. The yellow and green pie chart segments indicate the ratio of the number of cells at the two time points. (b) Distances between frequency vectors derived from three experimental rounds and the actual frequency vector for the PyMT-Cas9 tracing cell line, as labeled in Fig. 3c legend. (c) A magnified portion of Figure c displaying the results of experiment 3. Numbers 1-10 represent different proportions of specified active mutations ranging from 10% to 100%. The number 2 indicates a precise identification of the active mutations of hgRNA A21. (d) The transition diagram integrated with the learned transition probabilities for mutation alleles of hgRNA A21 in the PyMT-Cas9 tracing cell line. (e) The evolution of the frequencies of each mutation allele with the increase in the number of cell divisions.

We derived the corresponding transition diagram based on the mutation evolution network created for PyMT-Cas9. Following the experimental approach conducted on synthetic data, we conducted three rounds of experiments (Fig. 4b, 4c). In these experiments, we generated mutation evolution networks: one type that is reversible, another that is irreversible but differs from the mutation network we constructed, and a third one that has a structure completely matching our mutation evolution network but with incorrectly identified active mutation alleles. We then derived the corresponding transition diagrams for these networks and estimated the transition probabilities within the stochastic matrix using the maximum likelihood method.

The results demonstrated that the frequency vectors at the two time points obtained from these three rounds of experiments did not align with our experimental outcomes. Only when we used the mutation evolution network we created, correctly identified active mutation alleles, derived the appropriate transition diagram, and estimated the transition matrix through maximum likelihood, we can generate frequency vectors that matched our experimental results. The discrepancy between the learned frequency vector and the true frequency vector was < 0.02, measured using the Euclidean distance. This indicates that the difference between the learned and true values for each dimension in the frequency vector is less than 0.003.

Through an analysis of the learned transition probabilities, the likelihood of transitioning from the original hgRNA sequence to different mutations is not consistent (Fig. 4d). In the case of hgRNA A21, a 0.66 probability was found for remaining unchanged in each cell division, or in terms of the Markov model of transitioning to itself. This probability can be regarded as a measure of the transition rate for each hgRNA. Based on this transition rate, we can anticipate that the frequencies of various mutations in the case of A21 will reach a stable state after approximately ten cell divisions (Fig. 4e and Supplementary Fig. 5). This timeframe represents the scale over which hgRNA A21 can effectively track. It’s worth noting that we didn’t conduct the above analysis on hgRNA B25 because over 99% of the cells retained B25 unchanged.

Hence, there weren’t enough cells for the aforementioned analysis. This also aligns with the experimental findings that longer barcodes tend to have lower activity. From all the previous analyses, we have fully demonstrated the accuracy of the mutation evolution network and the correctness of the transition diagram of mutation alleles for hgRNA. They accurately depict the true evolutionary scenario of hgRNA as observed in our experiments.

### Reconstruct lineage relationships for individual cells using hgRNA

Based on our prior understanding of the previously discussed hgRNA characteristic, we evaluated its potential for reconstructing lineages at the single-cell level. We designed simulation experiments as follows: we began with a single cell, such as a fertilized egg, assuming that this cell already harbored *n* different hgRNAs with varying transition rates, specifically set to 0.5, 0.6, 0.7, 0.8, or 0.9. With each cell division, every hgRNA underwent mutation transitions based on a predetermined stochastic matrix. The cell cycle of each cell was simulated according to the characteristics of the PyMT-Cas9 tracing cell line. The simulation continued until specific time points were reached. The division history of each cell, along with mutations, was recorded, enabling us to construct real cell lineage relationships among all cells. Within the cells obtained, we calculated the progenitor distance between any two cells, defined as the average of the cell division times to their most recent common ancestor (Fig. 5a).

**Fig. 5:**
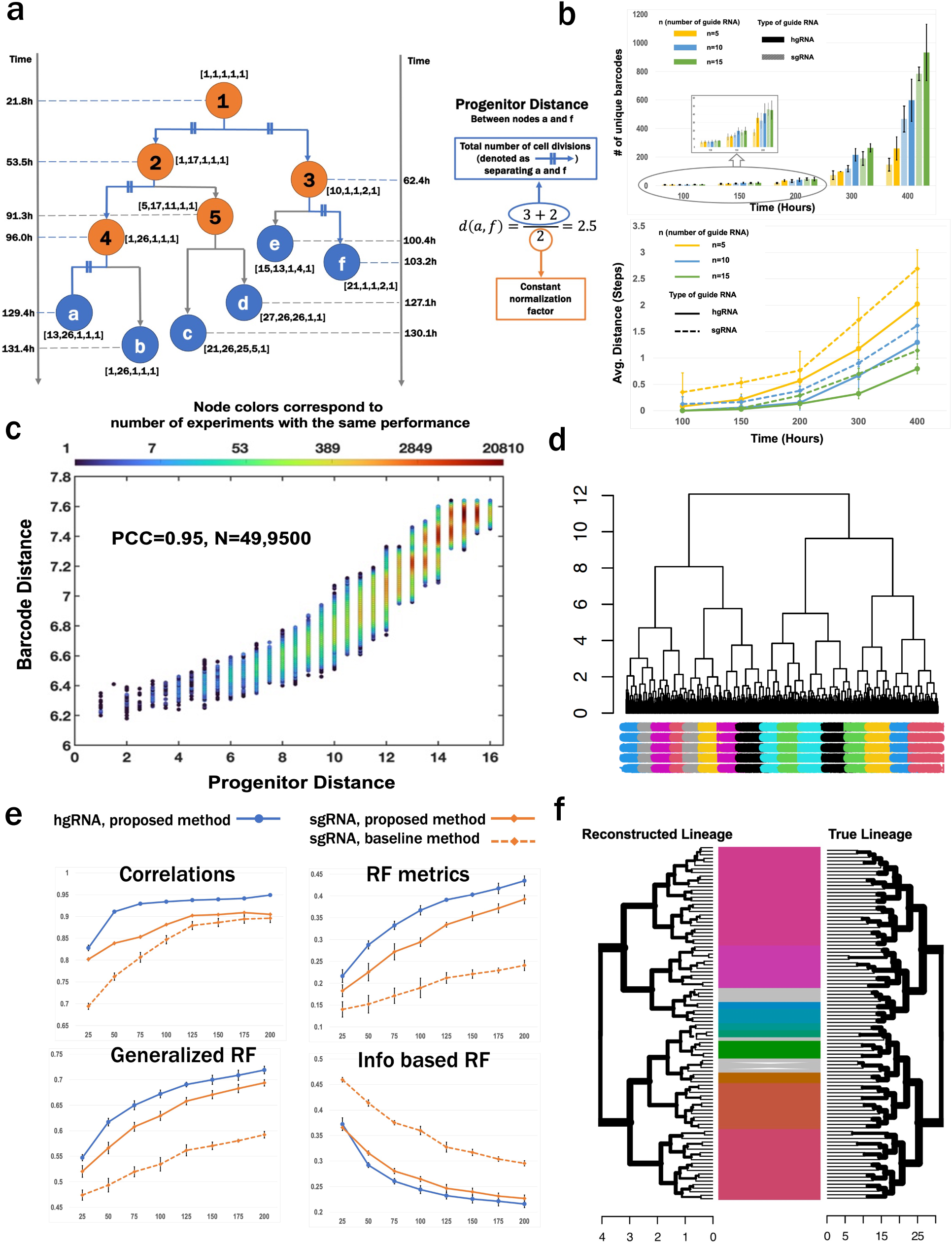
Reconstruction of cell lineage trees with simulated cell division using hgRNA. (a) In our simulation, an initial cell with 5 hgRNA undergoes several rounds of cell division, resulting in multiple cells. The blue leaf nodes denoted by letters, represent cell that truly existed at 100 hours, the yellow internal nodes, indicated by numbers, represent the ancestors of the leaf nodes. Each node is marked with a 5-dimensional vector representing the mutation alleles of each hgRNA within that cell, with number 1 indicating the original hgRNA sequence. The transition of mutation alleles for each hgRNA with each cell division is determined by a predesigned stochastic matrix. The time of each cell division is also recorded. (b) In the hgRNA system, each barcode shows higher compactness, indicating that cells sharing the same barcode are closer to each other compared to the sgRNA system. This observation remains consistent regardless of the number of guide RNAs used. (c) The Pearson correlation between pairwise progenitor distance and barcode distance for 1000 sampled cells from 53,512 simulated cells is 0.95, with 200 hgRNAs for recording. (d) Using barcode distances, a lineage tree for 1,000 sampled cells is reconstructed through hierarchical clustering. The descendants of the initial 16 cells resulting from the first four cell divisions are completely separated. Different colors represent one of the descendants of the 16 cells. (e) Comparing the accuracy of lineage reconstruction using different distance metrics, including their correlation with the progenitor distance, the Robinson-Foulds metric, a generalized Robinson-Foulds metric, and an information-based generalized Robinson-Foulds metric. For each parameter conditions, we calculate the mean accuracy and standard variant of 5 simulations. (f) Tree match between the true cell division history and reconstructed lineage trees for 100 sampled cells.

It is worth noting that canonical sgRNA is typically seen as a simplified version of the Markov model, where only the original sgRNA sequence itself can transition to any mutation, while all other mutations remain unchanged or transition only to themselves. Therefore, we created identical sgRNAs as designed for the hgRNA in these simulation experiments. Each sgRNA had the same number of mutation alleles as each hgRNA, and the transition probabilities from the original sgRNA to any mutation alleles were the same as those of hgRNA. The only difference is that all the other active mutation alleles in hgRNA were changed to inactive mutations for sgRNA. The simulation of cell divisions has also been recorded using sgRNA as we did with hgRNA. We planned to assess the differences in the effectiveness of two types of guide RNAs in reconstructing the lineage trees and evaluate the number of hgRNAs required to meet the demands of a specific biological question.

We first evaluated the issue of homoplasy for the two types of guide RNAs^18^. At each time point when we halted the cell division simulation, counted the number of unique barcodes for sgRNA and hgRNA. For each unique barcode, we calculated the progenitor distance between any pair of cells in which the unique barcode had been observed. The average progenitor distance among all pairs of cells with the same barcode was defined as the compactness of each mutation barcode. We aimed for the barcodes to be as compact as possible, signifying that cells bearing the same barcode were closely related and share very recent ancestors in their progenitor history, indicating similar biological characteristics. This strategy effectively alleviated the challenges posed by homoplasy.

*Our findings revealed that hgRNA outperformed sgRNA in generating a greater number of unique barcodes, and each of these barcodes exhibited higher compactness compared to sgRNA-generated barcodes*. This outcome suggests that hgRNA’s richer evolutionary pathways result in a broader spectrum of mutation combinations, ultimately leading to an increased diversity of unique barcodes. This increased diversity of mutation combinations enhances hgRNA’s ability to represent a broader range of cellular characteristics and lineage relationships, making it a valuable tool for lineage reconstruction and analysis (Fig. 5b).

As validated earlier, mutation alleles accumulate and become increasingly complex as cells undergo multiple rounds of divisions. Moreover, the compactness of barcodes indicates that cells sharing the same barcode are closely related. As a result, cells with significantly different barcodes should exhibit a considerable genetic distance in their progenitor histories.

Therefore, we defined the barcode distance between two cells based on their barcodes, utilizing the joint distribution probability of two mutation alleles during the Markov process to quantify the dissimilarity of two barcodes (Methods). The barcode distances and progenitor distances between cells are highly correlated with coefficient of 0.95 (Fig. 5c). Building upon this, we employed the barcode distances to reconstruct the dynamic process of cell division and applied hierarchical clustering algorithm to these barcode distances for lineage tree reconstruction.

To achieve that purpose, our simulation extended to 550 hours, generating a population of 53,512 cells. Within this group, 33 cells completed 15 cell divisions, 13,379 underwent 16, 37,192 achieved 17, and 2,908 reached 18 divisions. At the end of the simulation, we randomly sampled 1,000 cells for lineage relationship reconstruction, using hierarchical clustering informed by barcode distances, as depicted in Figure 5d. The barcode distances were also calculated based on the sgRNA system. To assess the performance of our cell distance definition compared to other approaches, we utilized another distance metric introduced for sgRNA^40^. Subsequently, we compared the lineage trees, reconstructed using different metrics, to the actual lineage tree to determine reconstruction fidelity. The accuracy of lineage reconstruction was determined using the Robinson-Foulds metric^41^, its generalized form^42^, and an information-based extension^43^.

The lineage trees reconstructed using our definition of barcode distance with the hgRNA system consistently outperformed those constructed with the sgRNA system. Notably, our hgRNA-based metric demonstrated consistent supremacy over the alternate sgRNA metric, as shown in Figure 5e. This superiority underscores our metric’s enhanced alignment with the actual number of cell divisions, thus more faithfully mirroring the cells’ developmental trajectories. Compared to sgRNA, the hgRNA system captures a more intricate evolutionary journey, gathering a richer trove of information on cellular division events. These findings are pivotal, as they illustrate that strategic selection of hgRNA sequences can lead to highly precise single-cell lineage tracing, as depicted in Figure 5f. *Our study confirms the practicality and effectiveness of using hgRNA for detailed single-cell lineage mapping*.

## Discussion

In this study, we created a PyMT-Cas9 tracing cell line with two hgRNA barcodes. These barcodes serve as markers to track genetic changes. We harvested cells from this tracing cell line at 5 days and 14 days to observe mutation alleles. We developed an advanced analytic pipeline and implemented that in LINEMAP to identify mutation alleles observed in each cell; therefore enabling us to calculate the frequency of occurrence for each mutation allele and provide quantitative information about the prevalence of different mutations.

Based on the observed mutation alleles and their frequencies, a mutation evolution network was constructed. This constructed network visually represents the evolutionary paths of mutation alleles, ranging in order of least complex to most complex forms. The evolutionary network provides a wealth of information about mutation evolution. For instance, the study assumes that mutation transitions are irreversible and must evolve according to the directions shown in the network. Furthermore, each mutation allele can only transition from one of its upstream alleles, and each active mutation allele can only transition to one of its downstream alleles. This directionality is a fundamental aspect of the model. The information derived from the mutation evolution network furnishes ample details for constructing transitions between mutations, forming the foundation for accurately describing how mutations transform and evolve over time.

The transition diagram built upon the mutation evolution network provides a graphical representation allele mutations’ progression. By analyzing mutation allele frequencies at two distinct time points, we can deduce the probabilities of transitions between these mutations. We tested the validity of the transition diagram and its associated probabilities through simulations of mutation networks across various scales. The crux of validation was to check if the mutation frequencies derived from the simulations, using our transition diagram and probabilities, matched the observed data at both time points. Our findings were affirmative; the alignment of simulation-derived mutation frequencies with empirical data was observed only when applying the authentic transition diagram and probabilities. Specifically, our results on the cell line data showed that only the transition diagram created based on our network could yield the mutation allele frequencies obtained in our experiments. This accuracy confirmation validates our network, confirming its efficacy in depicting the actual mutation transition dynamics.

By precisely characterizing the evolutionary path of hgRNA, LINEMAP can be used to reconstruct lineage relationships at the single-cell level based on hgRNA. There are several major methods for lineage reconstruction including distance, parsimony, likelihood, and Bayesian methods^18, 44^. In the context of CRISPR-Cas9 techniques, a variety of lineage reconstruction algorithms have been proposed, based on maximum parsimony^12,19,45–48^, maximum likelihood^23,24,49^, and distance-based methods^50,51^. Parsimony and maximum likelihood serve as fundamental principles for constructing a tree, whereas the internal node represent mutation events leading to the mutation state. However, these assumptions are not suitable for the hgRNA system, as previously demonstrated, due to the complexity and uncertainty in the evolution path of mutation alleles. Additionally, this type of method provides limited information about the true developmental timing of cell divisions. Existing distance-based methods often rely on heuristic approaches to define barcode distances, such as considering shared mutation events between two barcodes. However, these distance metrics are too crude to accurately capture the multiple rounds of cell divisions.

We defined a cell-cell distance based on the joint probability of their barcodes, representing the likelihood that two cells, resulting from the division of one cell, carry this specific barcode. Since the evolutionary paths of hgRNA have been fully characterized by the Markov process, this joint distribution can be computed. The lower the probability, the less likely these two barcodes occur simultaneously, indicating a greater distance between them. Importantly, based on our distance definition, even when two barcodes are identical, their distance is not zero. This is unique compared to previous distance definitions where cells sharing the same barcode have a distance of 0, when introducing the homoplasy issue.

The hgRNA system’s capacity for retargeting and persistent editing lends a complex nature to the evolution of its mutations, where a single mutation allele may arise from multiple evolutionary paths. Our experimental results confirm that by comprehensively understanding hgRNA’s evolutionary dynamics, the barcode distances we calculate are an accurate reflection mutation accumulation and indicates the number of cell divisions. Theoretically, as we increase the number of hgRNA sequences, the barcode distance will increasingly correlate with the progenitor distance, ultimately converging towards a perfect correlation. This will enable the complete reconstruction of a cell’s division history using our distance-based methodology. Our approach achieves unparalleled temporal and molecular precision, delineating each division at the single-cell level.

In conclusion, this study introduces a cutting-edge computational approach for detailed lineage reconstruction, enhancing our understanding of mutation allele evolution. The versatility of mutation evolution network and our distance-based lineage reconstruction method extend beyond to the hgRNA system. They can be used to characterize mutations across the genome and can be adapted to other sysems, including sgRNA and alternative tracing technologies. Research indicates that Cas9-mediated outcomes are context-independent, dictated solely by the target sequence, which allows us to predict the results in cell lines and anticipate them in primary cells^52–55^. Furthermore, LINEMAP could extend to other tracing strategies if the transition probabilities between states are computable^56^. Hence, the methodology and findings of this study could have wide-ranging implications, from understanding cell lineage in developmental biology to tracking the evolution of cancer cells. The specificity and detail provided by our study represent a valuable advancement for the scientific community.

## Data availability

## Code availability

## Acknowledgments

The authors acknowledge the funding support from the national Institutes of Health (grant number NIH R01CA244413 to D.G. and S.T.C.W., NIH R01CA251710, NIH U01CA253553, and NIH U01CA268813 to S.T.C.W). This research is also partially funded by T.T & W.F. Chao Foundation and John S. Dunn Research Foundation to S.T.C.W.

## Author contributions

S.T.C.W and D.G. conceived the idea of hgRNA evolution for single cell lineage tracing. L.W., W.D., Z.Y., D.G., and S.T.C.W. designed the study. S.T.C.W and D.G. funded and supervised the study. L.W., Z.Y. and J.S. conceived and designed the mathematical model, and L.W. wrote the code and performed the analyses. W.D. and D.G. designed and developed the tracing cell line. L.W., Z.Y., L.Y., S.Y.W., Z.W., X.Y. and K.H. designed the simulations and L.W. wrote the code and performed the analyses. L.W., Z.Y., J.S., C.F.E., S.Y.W., R.L.D. and S.T.C.W. performed bioinformatics analysis and evaluated the results. L.W., W.D., J.S., S.Y.W., R.L.D., C.F.E. and S.T.C.W wrote the first draft of the manuscript, and all authors approved the final draft.

## Methods

### Plasmids

Barcode A21 and B25 were selected from the barcode pool to generate MARC1 mouse line and cloned into pBA904 vector (Addgene#122238). For the barcode A21 plasmid, we replaced the selection marker Puro-T2A-BFP with Hygromycin-resistance gene. For the barcode B25 plasmid, we replaced the human promoter with mouse promoter as well as the selection marker gene by BFP. Both modifications were done using homologous recombination technology.

### Generating A21B25 barcode PyMT-Cas9 tracing cell line

A21B25 barcode PyMT cell line consist of three major parts, EMT tracing protein FSP fused with RFP-stop codon-GFP cassette, Cas9 protein, and A21B25 barcodes. PyMT EMT tracing cell line was constructed previously (PMID: 26560033). Briefly, FSP protein as an indicative protein for the EMT process was fused with RFP-stop codon-GFP cassette. When FSP protein is expressed, cells change florescence from RFP to GFP irreversibly which indicates the transition of epithelial state to mesenchymal state. Pseudotyped lentivirus containing Cas9, Puromycin-resistance gene, and GFP were added to PyMT EMT tracing cells, 2µg/ml puromycin were supplied to the cells 48h post infection for 5 days. Selected cells were used for adding A21 and B25 barcodes.

Plasmid containing barcode A21 or B25 were co-transfected with psPAX2 lentiviral packaging vector (Addgene# 12260) and pCMV-VSV-G plasmid (Addgene# 8454) into 293T cells using Lipo3000 according to manufacturer’s instruction. The supernatant was collected and titered 48h post transfection. PyMT-Cas9 cells were infected with Lentivirus containing A21 barcode at a MOI of 0.2-0.4 to ensure a single copy of the barcode in each cell. Selection medium supplemented with 600ug/ml hygromycin were added into the PyMT-Cas9 cells 48h post infection for 5 days. Selected PyMT-Cas9 cells containing A21 barcode were then infected with Lentivirus containing B25 barcode using the same MOI. The PyMT-Cas9 cells containing both A21 and B25 barcodes (BFP+) as well as in the epithelial states (RFP+) were then sorted and seeded as single clones in a 96-well plate. To generate the A21B25 barcode PyMT cell line, we picked out the single cell clones according to the following criteria: 1) all cells grown from the single cell clone are CFP positive; 2) there are both RFP+ and GFP+ cells in the clones and the ratio of RFP to GFP is between 20%-50%; and 3) the size of the clones is no less than the average clone size. 113 clones out of 1,200 total clones matched these criteria were selected for further verification. Genomic DNAs were isolated from all 113 clones as template for the quantitative amplification of A21 and B25 barcode using specific A21 and B25 barcode primers respectively. The clones having a A21 to B25 ratio between 0.9-1.1 were selected. Three clones out of 113 clones reached the final selection and were mixed at 1:1 ratio for the future experiments as A21B25 barcode PyMT cell line.

### Pipeline to identify the mutation alleles on single cell level

In our experiments, we utilized two hgRNA sequences, A21 and B25, with spacer sequences ‘GTTCCCGTCCAGTAATCGTG’ and ‘GTCGTTGTAGCAACCTATCGGGTG’, respectively.

#### 1. Filter the reads from the fastq files

Each read from the Read 2 (R2) FASTQ file is 90 bp lengths, contained 30 bp primer, and the 20 bp A21 hgRNA spacer sequence ‘*GTTCCCGTCCAGTAATCGTG’* in the example below, and PAM ‘*GGG’,* followed a part of the hgRNA scaffold. Each read from the Read 1 (R1) FASTQ file is 28 bp lengths, 16 bp cell barcode and 12 bp UMI are encoded. An example of the 28 bp read of Read 1 (R1) FASTQ file from CRISPR Screening: NGACCATTCTTAACAGGGTGATCGCTTT

An example of the 90 bp read of Read 2 (R2) FASTQ file from CRISPR Screening: aagcagtggtatcaacgcagagtacatgggGTTCCCGTCCAGTAATCGTGgggttagagctagaaatagcaagttaacctaaggctagtc

In the first step, for the purpose of reduce the unrelated reads in the following analysis, we applied a rough filter on the R2 reads by finding the longest sub-string of the hgRNA scaffold in each read. Reads that had longest sub-strings of the hgRNA scaffold shorter than 10 bases were considered too poor to be trusted for further analysis and were filtered out. Considering that the hgRNA scaffold should remain unchanged while the spacer sequence may get mutated during CRISPR-Cas9 edits, including at least a 10-base sub-string of the hgRNA scaffold in the reads is a very rough criterion.

#### 2. Determine the mutations using ampliCan

We used a R package called ‘ampliCan’ to identify the induced mutations^57^. The Insertions or deletions (indels) are identified by sequencing the targeted loci and comparing the sequenced reads to a reference sequence (original hgRNA like A21 we used here) using alignment algorithm. The ampliCan tool can accurately determine the true mutation efficiency. Each read in the Read 2 (R2) FASTQ file is aligned to A21 and B25, respectively, as we cannot determine which targeted loci have been sequenced in this read.

#### 3. Mutation description: construct a descriptive matrix for mutations

We used a descriptive matrix to structurally describe each specific mutation allele of genes (in this case, for hgRNA A21). As hgRNA A21 consists of 20 bp, with an additional 3 bp PAM, the total sequence of interest is 23 bp. Mutations such as deletions or mismatches can occur at each base, while insertions can occur between any two consecutive bases. Therefore, a matrix of size 45×5 has been constructed. The first 23 rows represent each base of the sequence, while the last 22 rows represent the ‘space’ between each two consecutive bases of the 23 bp sequence. The first column describes the mutation type, which could be either ‘mismatch’ or ‘deletion’ for the first 23 values, and ‘insertion’ for the last 22 values. The second column indicates the location of the mutation, with consecutive numbers from 1 to 23 for the bases and consecutive numbers from 1 to 22 for the spaces. The third column represents a number indicating the count of bases that have been changed or added. For the first 23 values, this number can only be 1 since we are considering single base mutations. However, for the last 22 values, which correspond to insertions, this number can be any positive integer as the length of the inserted sequence can vary theoretically. The fourth column contains the original bases, while the fifth column showcases the actual bases resulting from the mutation. Below, we provide a detail matrix describing a specific mutation for A21.

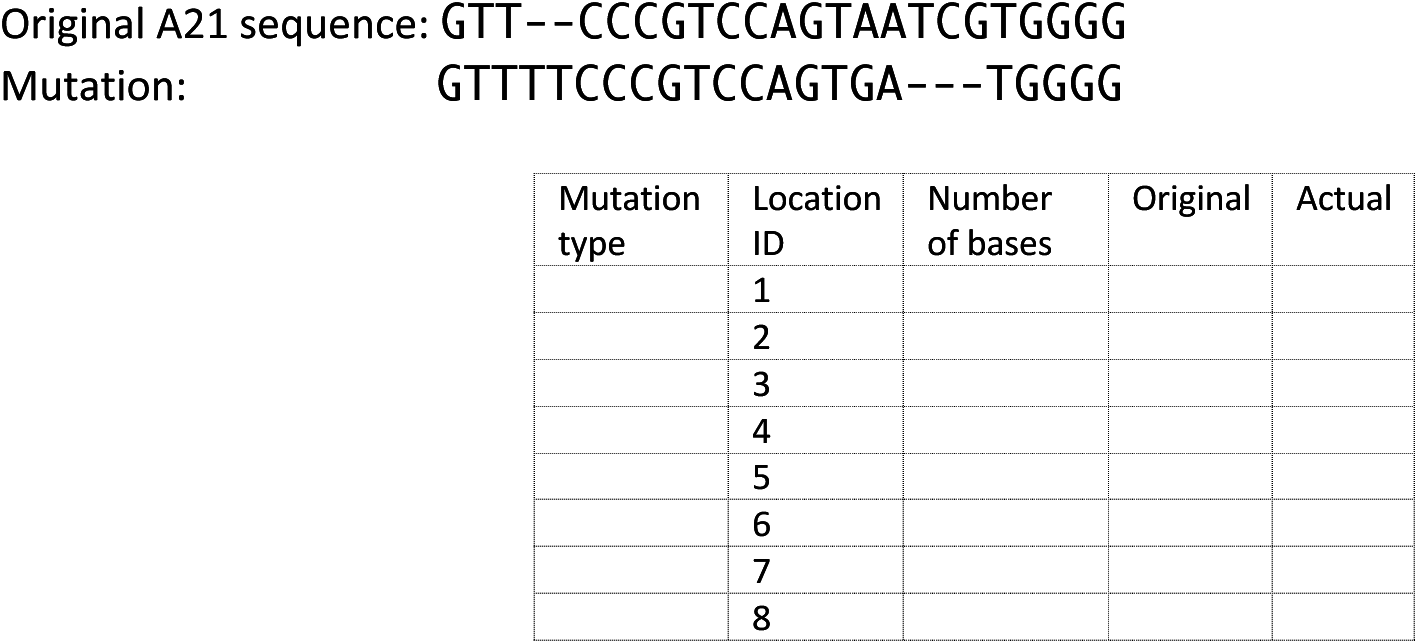
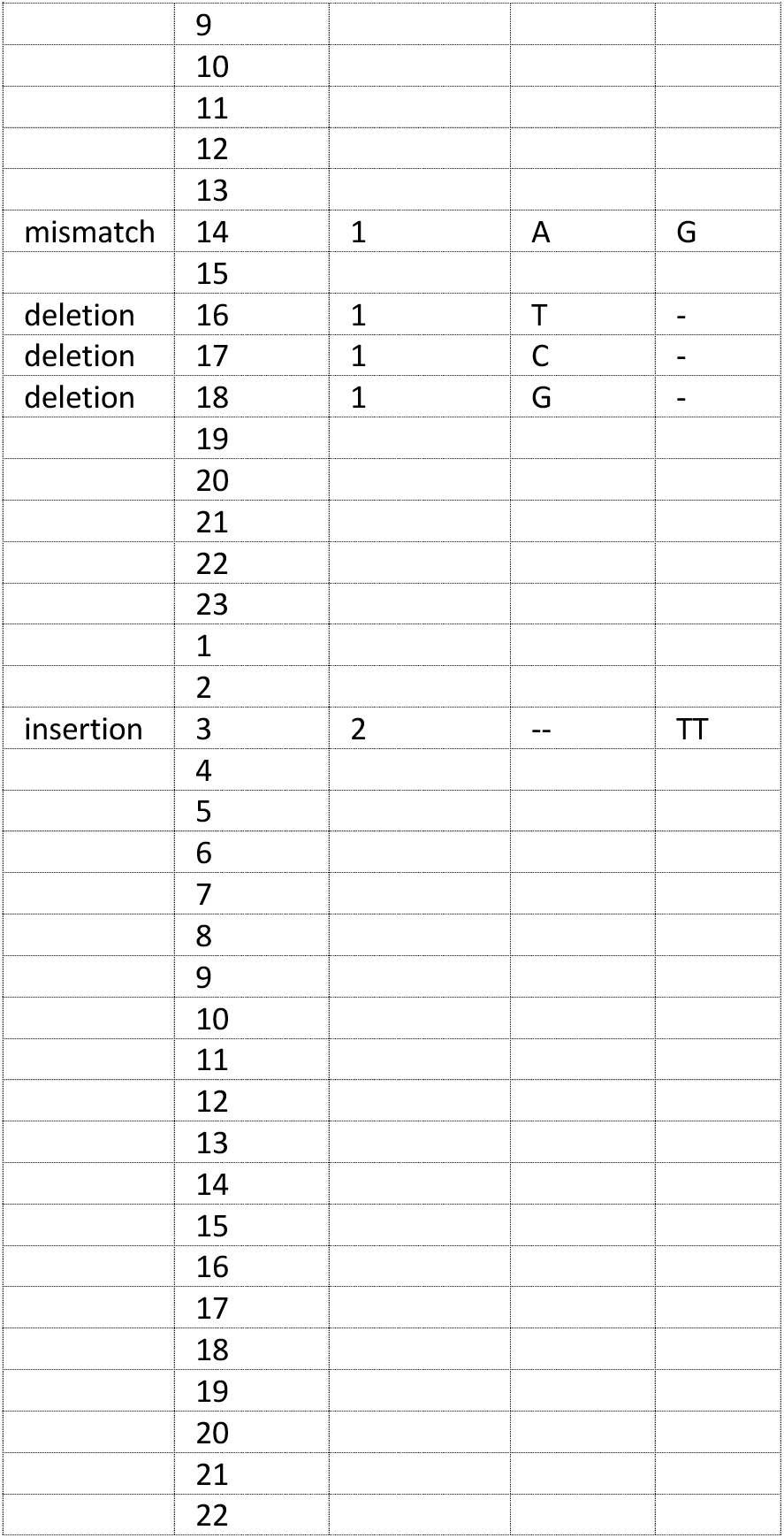

#### 4. Assign each read to A21 or B25

Each read will be aligned twice, once with hgRNA A21 as a reference and once with hgRNA B25 as a reference. After the mutations have been determined using the ampliCan tool, description matrices are constructed for A21 and B25 respectively. The third column of the descriptive matrix, which represents the ‘number of bases’ will be summed up to measure the complexity of the mutation compared to its original hgRNA sequence. The final complexity of the alignment will not only consider the complexity of the mutation on the hgRNA sequence, but also the alignment accuracy on the scaffold part, so we define a complexity score for the read as:

*Complexity score = length of scaffold - length of mismatch on scaffold – sum of number of bases*

The second filter will be applied to the scaffold part for quality check of the read. If the scaffold has a mismatch with the reference that exceeds 1/10 of the scaffold length, the alignment will be discarded for the corresponding reference hgRNA. Thus, each read will be finally assigned to either A21 or B25, or discarding them.

#### 5. Find the dominant mutation for each cell

Now we will take the Read 1 (R1) FASTQ file into consideration, which include the cell barcode information. Each cell may have tens or hundreds of reads, and among them, multiple mutations may have been identified. The dominant mutation for each cell will be determined based on the Complexity scores. The mutation with the highest Complexity score will be considered as the dominant mutation.

#### 6. Construct mutation evolution network

We constructed the Mutation Evolution Network to depict the progression of mutations. Essentially, the network is designed as a Directed Acyclic Graph (DAG). Every node within it represents a distinct mutation of the hgRNA. Each directed edge originates from a preceding mutation and leads to a subsequent mutation. In this process, the subsequent node encompasses all the locations mutated in its predecessor while introducing further mutated locations. The implementation of this concept utilizes the mutation description matrix. In the first column of the matrix, we assign a value of 0.5 for ‘mismatch’ mutations, 1 for ‘insertion’ and ‘deletion’ mutations, and 0 if there is no mutation at a particular location. The mutation vector of a child node should be greater than or equal to that of its parent node. If the mutation vectors are identical, then the third column of the description matrix of the child node should also be greater than or equal to that of its parent node. Moreover, there should be no other mutations occurring between the parent node and child node.

#### 7. Combine the networks of each sample into one

We have two samples collected at two different time points: 5 days and 14 days. Each sample will undergo steps 1 and 6 to create a mutation evolution network. Now, we will combine the two networks together. The mutation patterns from both samples will be merged, and a new mutation evolution description network will be constructed, following the process described in step 6. In the new network, each node will be presented as a pie chart, where the size of the pie chart is proportional to the percentage of occurrence of that mutation in each sample.

### Calculate the number of cell cycles at a time point

We simulated the process of cell division and check how many cell divisions would have happened when we sampled the cells at day 5 (120 hours after and) and day 14 (336 hours). As the PyMT-Cas9‘s double time or cell cycle is about 24 – 48 hours, and each cell is at the different stage of the cell cycle in time 0, we used a Truncated Normal Distribution to simulate the cell cycle time, with the truncated interval (24,48), and mean cell cycle time 36, and standard variation 5. For the first cycle, we used the uniform distribution to simulate the reminder of the cell cycle progression. We summed up the time of each cell cycle time and check how many cell cycles have completed. We did the simulation for 10,000 cells and calculated the frequency of the number of cell divisions.

### Maximum likelihood estimates by solving the nonlinear optimization problem

We demonstrated that the maximum likelihood estimates of the transition probabilities in the stochastic matrix *P* can be transformed into a nonlinear optimization problem.

Assume we have identified *n* mutation alleles (include the original hgRNA barcode). And there are *N*_1_ cells and *N*_2_ cells collected at time point 1 and time point 2, respectively. We represent the mutation alleles observed in each cell at time point 1 as 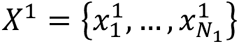, where *x_i_*, *i* ∈ {1, …, *N*_1_} is a one-hot vector indicating which mutation allele has been overserved in cell *i*. Similarly, at time point 2, we represent the mutation alleles observed in each cell as 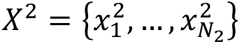. The frequency vector of the mutation alleles for time point 1 is denoted as 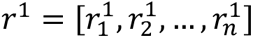, and for time point 2 as 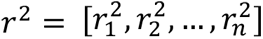. For each time point, we simulate the count of cell divisions, and we can calculate the frequency of each count of cell division. The frequency of each cell division count can be represented as 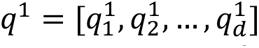 for time point 1, and 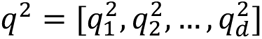 for time point 2, where *d* indicated the maximum count of cell division. Now, we can define the transition matrix for time point 1 and time point 2 as follows: 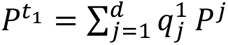 and 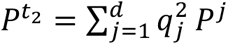, where *P* represents the stochastic matrix of the Markov model the mutation transition obeys at each cell division, and *P^j^* is the *j*th power of matrix *P* represents the transition matrix of mutations after a specific count of cell division *j*.

At time point 1, given samples, 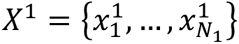, we seek the transition matrix *P*^*t*_1_^ that maximizes the log likelihood function defined as follows:

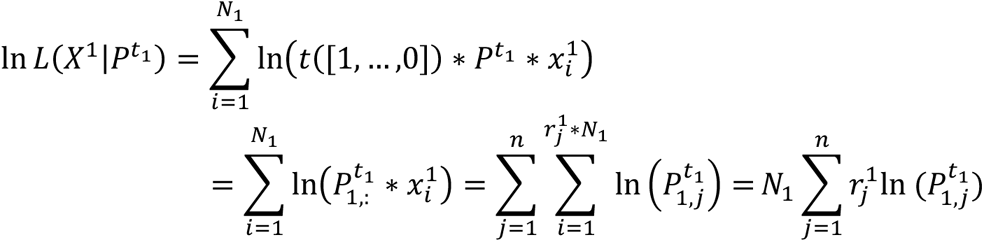

Here the vector [1, …, 0] represents the frequency vector at the start time when all the hgRNA barcodes are intact. 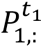 reprensents the frist row of transition matrix *P*^*t*_1_^, and 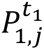 represents the element in the first row and *j*th column of transition matrix *P*^*t*_1_^.

Using the Lagrange multiplier method, we can obtain the maximum likelihood estimate of 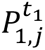 as:

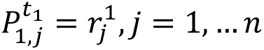

Or in matrix product form:

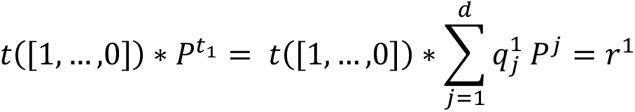

The same approach applies at time point 2:

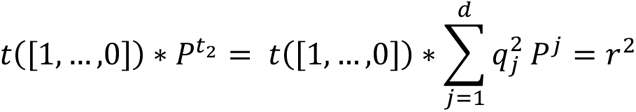

So to compute the maximum likelihood estimate of each transition probability in the stochastic matrix *P* of the Markov Model, we transferred the problem to a nonlinear optimization problem defined as follows:

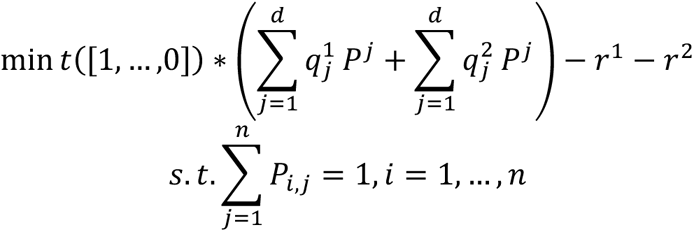

This nonlinear optimization problem was solved using the ‘parcma’ R package^58^, which provides function to solve system of nonlinear equations. The package uses the Gauss-Newton optimization algorithm to optimize the objective function with tolerance set to be 0.0001.

### Calculate the cell-cell distance based on the hgRNA barcodes

Let’s assume we have *N* samples, each containing *m* hgRNAs. Each hgRNA has the same number of mutation alleles, denoted as *n*. We represent the barcode of each cell as a vector [*l*_1_, …, *l_m_* where *l_k_* is a number indicating the mutation allele for hgRNA *k*, *k* ∈ {1,2, …, *m*}. At a specific time point *t*, the transition matrix of hgRNA *k* is defined as previous mentioned and is denoted as 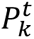. Assume cell *i* has barcode 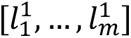 and cell *j* has barcode 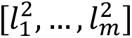, so the joint probability of cells *i* and *j* originating from one cell division, carrying these two barcodes can be calculated as:

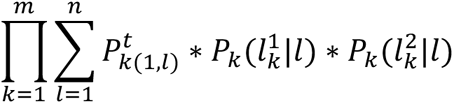

where 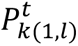 denotes the element in the first row and *l*th column of transition matrix 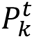, indicating the frequency of mutation allele *l* in the *k*th hgRNA, and 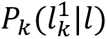 and 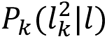 denote the probability of mutation allele *l* transit to mutation 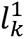 and 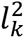 with one cell division.

To convert probabilities into distances, we take the natural logarithm of the inverse of the probabilities, and then define the distance between cell *i* and *j* as:

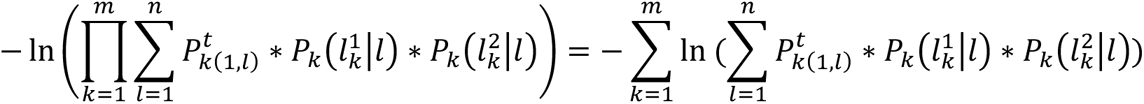

### Tree accuracy estimation

We used the Robinson–Foulds distance to calculate the distance between two lineage trees. This measure counts the number of bipartition splits that occur in both trees. The Generalized Robinson– Foulds metric matches splits in one tree with similar splits in the other, and each pair is assigned a similarity score. The sum of these scores quantifies the similarity between the two trees. The Information-based Generalized Robinson–Foulds metric introduced a probabilistic measure of split similarity. The calculation of all these metrics is implemented using the ‘TreeDist’ R package^59^. The tanglegram for tree matching is implemented using the ‘dendextend’ R package^60^.

## Reference

[1] Li, Z., Yang, W., Wu, P., Shan, Y., Zhang, X., Chen, F., Yang, J., & Yang, J. R. Reconstructing cell lineage trees with genomic barcoding: Approaches and applications. Journal of genetics and genomics = Yi chuan xue bao, S1673–8527(23)00121-2 (2023). Advance online publication. 10.1016/j.jgg.2023.05.011

[2] Gabbutt, C., Schenck, R.O., Weisenberger, D.J. et al. Fluctuating methylation clocks for cell lineage tracing at high temporal resolution in human tissues. Nat Biotechnol 40, 720–730 (2022). 10.1038/s41587-021-01109-w

[3] Wu, S. S., Lee, J. H., & Koo, B. K. Lineage Tracing: Computational Reconstruction Goes Beyond the Limit of Imaging. Molecules and cells, 42(2), 104–112 (2019). 10.14348/molcells.2019.0006

[4] Zhang, Y., Zeng, F., Han, X. et al. Lineage tracing: technology tool for exploring the development, regeneration, and disease of the digestive system. Stem Cell Res Ther 11, 438 (2020). 10.1186/s13287-020-01941-y

[5] Jia, X., Shen, G., Jia, J., Zhang, Y., Zhang, D., Li, W., Zhang, J., Huang, X., & Tian, J. Lineage Tracing and Molecular Real-Time Imaging of Cancer Stem Cells. Biosensors, 12(9), 703 (2022). 10.3390/bios12090703

[6] Wang, L. L., Serrano, C., Zhong, X., Ma, S., Zou, Y., & Zhang, C. L. Revisiting astrocyte to neuron conversion with lineage tracing in vivo. Cell, 184(21), 5465–5481 (2021).

[7] Gabbutt, C., Wright, N. A., Baker, A. M., Shibata, D., & Graham, T. A. Lineage tracing in human tissues. The Journal of Pathology, 257(4), 501–512 (2022).

[8] Abyzov, A., & Vaccarino, F. M. Cell lineage tracing and cellular diversity in humans. Annual Review of Genomics and Human Genetics, 21, 101–116 (2020).

[9] Weinreb, C., Rodriguez-Fraticelli, A., Camargo, F. D., & Klein, A. M. Lineage tracing on transcriptional landscapes links state to fate during differentiation. Science, 367(6479), eaaw3381 (2020).

[10] Zhou, L., Ken, H. O., Wong, T. L., Zhang, Z., Chan, C. H., Loong, J. H., … & Ma, S. Lineage tracing and single-cell analysis reveal proliferative Prom1+ tumour-propagating cells and their dynamic cellular transition during liver cancer progression. Gut, 71(8), 1656–1668 (2022).

[11] Jinek, M., Chylinski, K., Fonfara, I., Hauer, M., Doudna, J. A., & Charpentier, E. A programmable dual-RNA–guided DNA endonuclease in adaptive bacterial immunity. science, 337(6096), 816–821 (2012).

[12] Spanjaard, B., Hu, B., Mitic, N. et al. Simultaneous lineage tracing and cell-type identification using CRISPR–Cas9-induced genetic scars. Nat Biotechnol 36, 469–473 (2018). 10.1038/nbt.4124

[13] Michlits, G., Hubmann, M., Wu, SH. et al. CRISPR-UMI: single-cell lineage tracing of pooled CRISPR–Cas9 screens. Nat Methods 14, 1191–1197 (2017). 10.1038/nmeth.4466

[14] Gardner, A., Morgan, D., Al’Khafaji, A., & Brock, A. Functionalized Lineage Tracing for the Study and Manipulation of Heterogeneous Cell Populations. Methods in molecular biology (Clifton, N.J.), 2394, 109–131 (2022). 10.1007/978-1-0716-1811-0_8

[15] Boettcher, M., Covarrubias, S., Biton, A., Blau, J., Wang, H., Zaitlen, N., & McManus, M. T. Tracing cellular heterogeneity in pooled genetic screens via multi-level barcoding. BMC genomics, 20(1), 1–9 (2019).

[16] Wagner, D. E., & Klein, A. M. Lineage tracing meets single-cell omics: opportunities and challenges. Nature Reviews Genetics, 21(7), 410–427 (2020).

[17] Fang, W., Bell, C. M., Sapirstein, A., Asami, S., Leeper, K., Zack, D. J., … & Kalhor, R. Quantitative fate mapping: A general framework for analyzing progenitor state dynamics via retrospective lineage barcoding. Cell, 185(24), 4604–4620 (2022).

[18] McKenna, A., & Gagnon, J. A. Recording development with single cell dynamic lineage tracing. Development, 146(12), dev169730 (2019). 10.1242/dev.169730

[19] McKenna, A., Findlay, G. M., Gagnon, J. A., Horwitz, M. S., Schier, A. F., & Shendure, J. Whole-organism lineage tracing by combinatorial and cumulative genome editing. Science (New York, N.Y.), 353(6298), aaf7907 (2016). 10.1126/science.aaf7907

[20] Kester, L., & van Oudenaarden, A. Single-Cell Transcriptomics Meets Lineage Tracing. Cell stem cell, 23(2), 166–179 (2018). 10.1016/j.stem.2018.04.014

[21] Raj, B., Wagner, D., McKenna, A. et al. Simultaneous single-cell profiling of lineages and cell types in the vertebrate brain. Nat Biotechnol 36, 442–450 (2018). 10.1038/nbt.4103

[22] A., Florescu, M., Baron, C., et al. Whole-organism clone tracing using single-cell sequencing. Nature 556, 108–112 (2018). 10.1038/nature25969

[23] Bowling, S., Sritharan, D., Osorio, F. G., Nguyen, M., Cheung, P., Rodriguez-Fraticelli, A., … & Camargo, F. D. An engineered CRISPR-Cas9 mouse line for simultaneous readout of lineage histories and gene expression profiles in single cells. Cell, 181(6), 1410–1422 (2020).

[24] Chan, M. M., Smith, Z. D., Grosswendt, S., Kretzmer, H., Norman, T. M., Adamson, B., … & Weissman, J. S. Molecular recording of mammalian embryogenesis. Nature, 570(7759), 77–82 (2019).

[25] Quinn, J. J., Jones, M. G., Okimoto, R. A., Nanjo, S., Chan, M. M., Yosef, N., … & Weissman, J. S. Single-cell lineages reveal the rates, routes, and drivers of metastasis in cancer xenografts. Science (New York, N.Y.), 371(6532), eabc1944 (2021).

[26] Simeonov, K. P., Byrns, C. N., Clark, M. L., Norgard, R. J., Martin, B., Stanger, B. Z., Shendure, J., McKenna, A., & Lengner, C. J. Single-cell lineage tracing of metastatic cancer reveals selection of hybrid EMT states. Cancer cell, 39(8), 1150–1162.e9 (2021). 10.1016/j.ccell.2021.05.005

[27] Yang, D., Jones, M. G., Naranjo, S., Rideout, W. M., Min, K. H. J., Ho, R., … & Weissman, J. S. Lineage tracing reveals the phylodynamics, plasticity, and paths of tumor evolution. Cell, 185(11), 1905–1923 (2022).

[28] Paquet, D., Kwart, D., Chen, A. et al. Efficient introduction of specific homozygous and heterozygous mutations using CRISPR/Cas9. Nature 533, 125–129 (2016). 10.1038/nature17664

[29] Lieber, M. R. The mechanism of double-strand DNA break repair by the nonhomologous DNA end-joining pathway. Annual review of biochemistry, 79, 181–211 (2010).

[30] Shen, M.W., Arbab, M., Hsu, J.Y. et al. Predictable and precise template-free CRISPR editing of pathogenic variants. Nature 563, 646–651 (2018). 10.1038/s41586-018-0686-x

[31] Perli, S. D., Cui, C. H., & Lu, T. K. Continuous genetic recording with self-targeting CRISPR-Cas in human cells. Science, 353(6304), aag0511 (2016).

[32] Kalhor, R., Mali, P., & Church, G. M. Rapidly evolving homing CRISPR barcodes. Nature methods, 14(2), 195–200 (2017). 10.1038/nmeth.4108

[33] Kalhor, R., Kalhor, K., Mejia, L., Leeper, K., Graveline, A., Mali, P., & Church, G. M. Developmental barcoding of whole mouse via homing CRISPR. Science (New York, N.Y.), 361(6405), eaat9804 (2018). 10.1126/science.aat9804

[34] Leeper, K., Kalhor, K., Vernet, A. et al. Lineage barcoding in mice with homing CRISPR. Nat Protoc 16, 2088–2108 (2021). 10.1038/s41596-020-00485-y

[35] Zhang, W., Bado, I. L., Hu, J., Wan, Y. W., Wu, L., Wang, H., Gao, Y., Jeong, H. H., Xu, Z., Hao, X., Lege, B. M., Al-Ouran, R., Li, L., Li, J., Yu, L., Singh, S., Lo, H. C., Niu, M., Liu, J., Jiang, W., … Zhang, X. H. The bone microenvironment invigorates metastatic seeds for further dissemination. Cell, 184(9), 2471–2486.e20 (2021). 10.1016/j.cell.2021.03.011

[36] Bado, I. L., Zhang, W., Hu, J., Xu, Z., Wang, H., Sarkar, P., … & Zhang, X. H. F. The bone microenvironment increases phenotypic plasticity of ER+ breast cancer cells. Developmental cell, 56(8), 1100–1117 (2021).

[37] Lv, J., Wu, S., Wei, R., Li, Y., Jin, J., Mu, Y., Zhang, Y., Kong, Q., Weng, X., & Liu, Z. The length of guide RNA and target DNA heteroduplex effects on CRISPR/Cas9 mediated genome editing efficiency in porcine cells. Journal of veterinary science, 20(3), e23 (2019). 10.4142/jvs.2019.20.e23

[38] Zhang, J. P., Li, X. L., Neises, A., Chen, W., Hu, L. P., Ji, G. Z., Yu, J. Y., Xu, J., Yuan, W. P., Cheng, T., & Zhang, X. B. Different Effects of sgRNA Length on CRISPR-mediated Gene Knockout Efficiency. Scientific reports, 6, 28566 (2016). 10.1038/srep28566

[39] Cvijović, I., Good, B. H., Jerison, E. R., & Desai, M. M. Fate of a mutation in a fluctuating environment. Proceedings of the National Academy of Sciences of the United States of America, 112(36), E5021–E5028 (2015). 10.1073/pnas.1505406112

[40] Salvador-Martínez, I., Grillo, M., Averof, M., & Telford, M. J. Is it possible to reconstruct an accurate cell lineage using CRISPR recorders?. eLife, 8, e40292 (2019). 10.7554/eLife.40292

[41] Robinson, D. F., & Foulds, L. R. Comparison of phylogenetic trees. Mathematical biosciences, 53(1-2), 131–147 (1981).

[42] Nye, T. M., Lio, P., & Gilks, W. R. A novel algorithm and web-based tool for comparing two alternative phylogenetic trees. Bioinformatics, 22(1), 117–119 (2006).

[43] Smith, M. R. (2020). Information theoretic generalized Robinson–Foulds metrics for comparing phylogenetic trees. Bioinformatics, 36(20), 5007–5013 (2020).

[44] Yang, Z., Rannala, B. Molecular phylogenetics: principles and practice. Nat Rev Genet 13, 303–314 (2012). 10.1038/nrg3186

[45] Raj, B., Gagnon, J. A., & Schier, A. F. Large-scale reconstruction of cell lineages using single-cell readout of transcriptomes and CRISPR-Cas9 barcodes by scGESTALT. Nature protocols, 13(11), 2685–2713 (2018). 10.1038/s41596-018-0058-x

[46] Alemany, A., Florescu, M., Baron, C. et al. Whole-organism clone tracing using single-cell sequencing. Nature 556, 108–112 (2018). 10.1038/nature25969

[47] Jones, M.G., Khodaverdian, A., Quinn, J.J. et al. Inference of single-cell phylogenies from lineage tracing data using Cassiopeia. Genome Biol 21, 92 (2020). 10.1186/s13059-020-02000-8

[48] Feng, J., Dewitt III, W. S., McKenna, A., Simon, N., Willis, A. D., & Matsen IV, F. A. Estimation of cell lineage trees by maximum-likelihood phylogenetics. The annals of applied statistics, 15(1), 343 (2021).

[49] Zafar, H., Lin, C., & Bar-Joseph, Z. (2020). Single-571 cell lineage tracing by integrating CRISPR-Cas9 mutations with transcriptomic data. Nature communications, 11(1), 1–14.

[50] Hwang, B., Lee, W., Yum, SY. et al. Lineage tracing using a Cas9-deaminase barcoding system targeting endogenous L1 elements. Nat Commun 10, 1234 (2019). 10.1038/s41467-019-09203-z

[51] Frieda, K., Linton, J., Hormoz, S. et al. Synthetic recording and *in situ* readout of lineage information in single cells. Nature 541, 107–111 (2017). 10.1038/nature20777

[52] Allen, F., Crepaldi, L., Alsinet, C. et al. Predicting the mutations generated by repair of Cas9-induced double-strand breaks. Nat Biotechnol 37, 64–72 (2019). 10.1038/nbt.4317

[53] Hsu, P., Scott, D., Weinstein, J. et al. DNA targeting specificity of RNA-guided Cas9 nucleases. Nat Biotechnol 31, 827–832 (2013). 10.1038/nbt.2647

[54] van Overbeek, M., Capurso, D., Carter, M. M., Thompson, M. S., Frias, E., Russ, C., Reece-Hoyes, J. S., Nye, C., Gradia, S., Vidal, B., Zheng, J., Hoffman, G. R., Fuller, C. K., & May, A. P. DNA Repair Profiling Reveals Nonrandom Outcomes at Cas9-Mediated Breaks. Molecular cell, 63(4), 633–646 (2016). 10.1016/j.molcel.2016.06.037

[55] Chen, W., McKenna, A., Schreiber, J., Haeussler, M., Yin, Y., Agarwal, V., … & Shendure, J. Massively parallel profiling and predictive modeling of the outcomes of CRISPR/Cas9-mediated double-strand break repair. Nucleic acids research, 47(15), 7989–8003 (2019).

[56] Pei, W., Feyerabend, T., Rössler, J., et al. *Polylox* barcoding reveals haematopoietic stem cell fates realized *in vivo*. Nature 548, 456–460 (2017). 10.1038/nature23653

## Reference

[57] Labun, K., Guo, X., Chavez, A., Church, G., Gagnon, J. A., & Valen, E. Accurate analysis of genuine CRISPR editing events with ampliCan. Genome research, 29(5), 843–847 (2019). 10.1101/gr.244293.118

[58] Borchers, H. W. (2023). pracma: Practical Numerical Math Functions (Version 2.4.2). Available from https://cran.r-project.org/package=pracma

[59] Smith MR (2020). TreeDist: Distances between Phylogenetic Trees. R package version 2.6.3. doi:10.5281/zenodo.3528124.

[60] Galili T (2015). “dendextend: an R package for visualizing, adjusting, and comparing trees of hierarchical clustering.”

