## supplemental material for "Charting Single Cell Lineage Dynamics and Mutation Networks via Homing CRISPR"

**Supplementary Figures and Tables**

Fig. 1. The Sankey diagram displays the frequencies of different sequences inserted at various locations.

Fig. 2. The experimental settings were identical to those described in the main Fig. 3, except that 10% of nodes were designated as active mutations in the simulated data. The number 1 denotes a perfect match between specified active mutations and those utilized in the actual simulated data, at 10%.

Fig. 3. The experimental settings were identical to those described in the main Fig. 3, except that 30% of nodes were designated as active mutations in the simulated data. The number 3 denotes a perfect match between specified active mutations and those utilized in the actual simulated data, at 30%.

Fig. 4. The experimental settings were identical to those described in the main Fig. 3, except that 40% of nodes were designated as active mutations in the simulated data. The number 4 denotes a perfect match between specified active mutations and those utilized in the actual simulated data, at 40%.

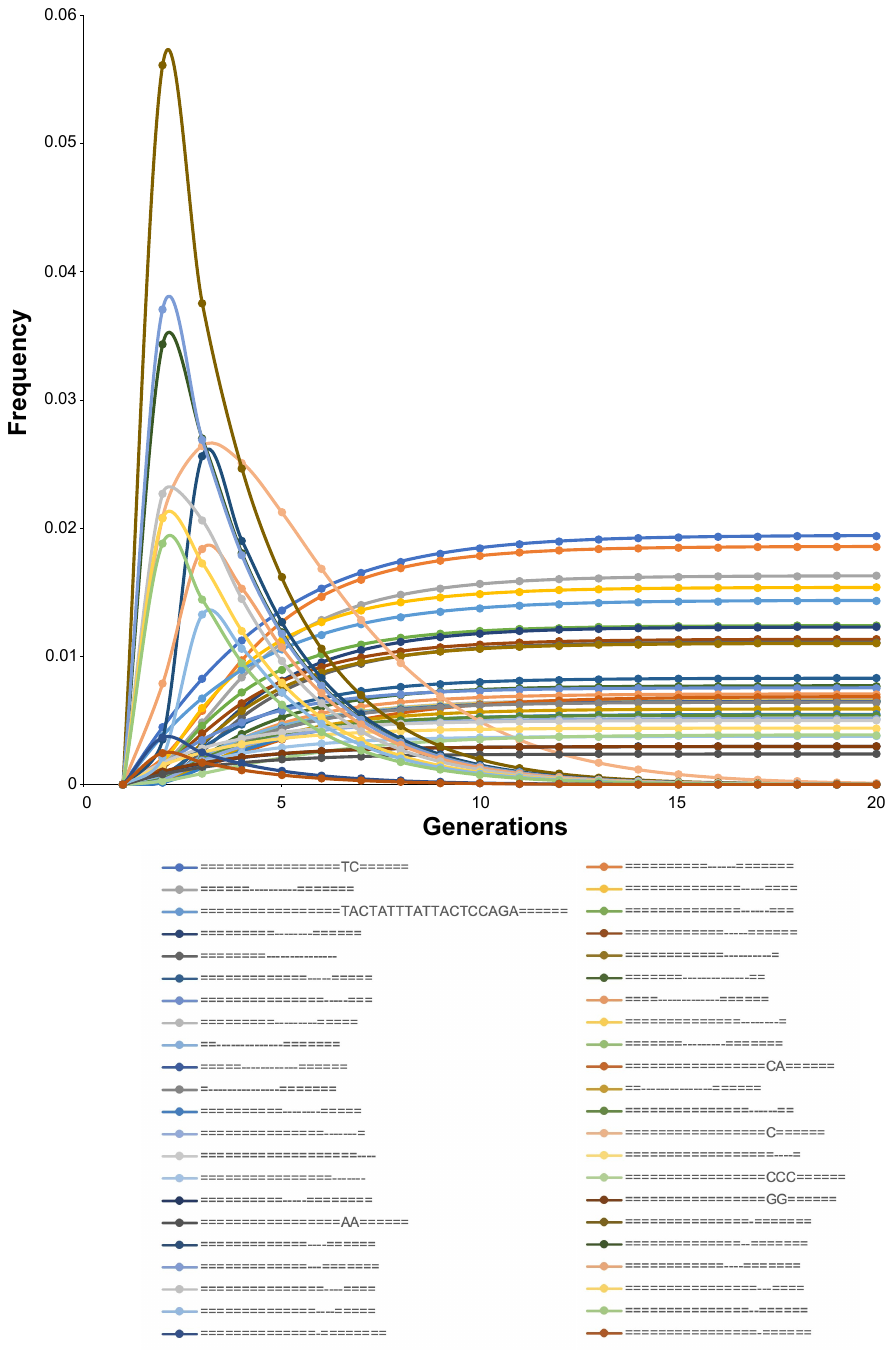

Fig. 5. The evolution of the frequencies of mutation alleles, not depicted in Fig. 4e, with the increase in the number of cell divisions.

**Table 1.** Characteristic of mutation alleles for hgRNA A21 observed at day 5 and day 14. '-' denotes deleted base pairs, letters indicate inserted base sequences, and '=' represents non-mutated base pairs.

| Mutation used in the mutation evolution network | No. cells at day 5 | No. cells at day 14 | Total No. cells | Frequency at day 5 | Frequency at day 14 | Active mutation (1) or not (0) |
| --- | --- | --- | --- | --- | --- | --- |
| =============------==== | 628 | 1372 | 2000 | 0.2583 | 0.4385 | 0 |
| ======================= | 599 | 84 | 683 | 0.2464 | 0.02685 | 1 |
| =============---------- | 109 | 312 | 421 | 0.04484 | 0.09971 | 0 |
| =================CC====== | 108 | 229 | 337 | 0.04443 | 0.07319 | 0 |
| ==-----------------==== | 72 | 86 | 158 | 0.02962 | 0.02748 | 0 |
| ===========-----======= | 49 | 48 | 97 | 0.02016 | 0.01534 | 0 |
| ==========-------====== | 33 | 60 | 93 | 0.01357 | 0.01918 | 0 |
| ============--------=== | 35 | 57 | 92 | 0.01440 | 0.01822 | 0 |
| =================TC====== | 35 | 53 | 88 | 0.01440 | 0.01694 | 0 |
| ========-----------==== | 22 | 64 | 86 | 0.009050 | 0.02045 | 0 |
| ============---------== | 22 | 63 | 85 | 0.009050 | 0.02013 | 0 |
| ==========------======= | 33 | 50 | 83 | 0.01357 | 0.01598 | 0 |
| ======----------======= | 31 | 42 | 73 | 0.01275 | 0.01342 | 0 |
| ==============-----==== | 26 | 46 | 72 | 0.01070 | 0.01470 | 0 |
| =================C====== | 62 | 8 | 70 | 0.02550 | 0.002557 | 1 |
| ==============------=== | 29 | 30 | 59 | 0.01193 | 0.009588 | 0 |
| ============----======= | 47 | 8 | 55 | 0.01933 | 0.002557 | 1 |
| ============-----====== | 22 | 30 | 52 | 0.009050 | 0.009588 | 0 |
| =========--------====== | 16 | 36 | 52 | 0.006582 | 0.01151 | 0 |
| ===============-======= | 46 | 5 | 51 | 0.01892 | 0.001598 | 1 |
| ============----------= | 19 | 31 | 50 | 0.007816 | 0.009907 | 0 |
| =============----====== | 27 | 20 | 47 | 0.01111 | 0.006392 | 1 |
| ========--------------- | 10 | 35 | 45 | 0.004114 | 0.01119 | 0 |
| ==============--======= | 33 | 9 | 42 | 0.01357 | 0.002876 | 1 |
| =============---======= | 36 | 4 | 40 | 0.01481 | 0.001278 | 1 |
| =============-----===== | 7 | 31 | 38 | 0.002879 | 0.009907 | 0 |
| ===============-----=== | 11 | 25 | 36 | 0.004525 | 0.007990 | 0 |
| ==============--------= | 18 | 13 | 31 | 0.007404 | 0.004155 | 0 |
| ==--------------======= | 16 | 14 | 30 | 0.006582 | 0.004474 | 0 |
| ====-------------====== | 13 | 16 | 29 | 0.005348 | 0.005113 | 0 |
| ===============----==== | 22 | 6 | 28 | 0.009050 | 0.001918 | 1 |
| =======---------======= | 14 | 14 | 28 | 0.005759 | 0.004474 | 0 |
| =---------------======= | 12 | 15 | 27 | 0.004936 | 0.004794 | 0 |
| ===================---- | 13 | 12 | 25 | 0.005348 | 0.003835 | 0 |
| ===============------== | 8 | 17 | 25 | 0.003291 | 0.005433 | 0 |
| ===============--====== | 17 | 7 | 24 | 0.006993 | 0.002237 | 1 |
| ===============-------= | 10 | 14 | 24 | 0.004114 | 0.004474 | 0 |
| =====------------====== | 6 | 18 | 24 | 0.002468 | 0.005753 | 0 |
| =================CA====== | 2 | 21 | 23 | 0.0008227 | 0.006711 | 0 |
| ================---==== | 19 | 3 | 22 | 0.007816 | 0.0009588 | 1 |
| ==================----= | 4 | 18 | 22 | 0.001645 | 0.005753 | 0 |
| ==---------------====== | 5 | 16 | 21 | 0.002057 | 0.005113 | 0 |
| =======--------------== | 9 | 10 | 19 | 0.003702 | 0.003196 | 0 |
| ==============----===== | 17 | 1 | 18 | 0.006993 | 0.0003196 | 1 |
| ================------- | 3 | 15 | 18 | 0.001234 | 0.004794 | 0 |
| =================GG====== | 4 | 11 | 15 | 0.001645 | 0.003516 | 0 |
| ==========--------===== | 1 | 16 | 17 | 0.0004113 | 0.005113 | 0 |
| =========---------===== | 10 | 5 | 15 | 0.004114 | 0.001598 | 0 |
| ================-====== | 11 | 3 | 14 | 0.004525 | 0.0009588 | 1 |
| =================AA====== | 0 | 12 | 12 | 0 | 0.003835 | 0 |
| =================CCC====== | 0 | 12 | 12 | 0 | 0.003835 | 0 |
| ==========-----======== | 10 | 2 | 12 | 0.004114 | 0.0006392 | 0 |
| ==============-======== | 10 | 0 | 10 | 0.004114 | 0 | 1 |
| =================TACTATTTATTACTCCAGA====== | 10 | 0 | 10 | 0.004114 | 0 | 0 |
| Disregarded mutations | No. cells at day 5 | No. cells at day 14 | Total No. cells |  |  |  |
| =============-------=== | 9 | 9 | 18 |  |  |  |
| ==================--=== | 7 | 9 | 16 |  |  |  |
| ==============-------== | 9 | 7 | 16 |  |  |  |
| ================AAAAGCTCCG======= | 6 | 9 | 15 |  |  |  |
| ==============--------- | 5 | 9 | 14 |  |  |  |
| ===------------======== | 7 | 6 | 13 |  |  |  |
| ================------= | 3 | 9 | 12 |  |  |  |
| ===========-------===== | 6 | 6 | 12 |  |  |  |
| ==========------------= | 4 | 8 | 12 |  |  |  |
| =================G====== | 7 | 4 | 11 |  |  |  |
| ==============---====== | 8 | 3 | 11 |  |  |  |
| ==========---------==== | 7 | 4 | 11 |  |  |  |
| =================CCA====== | 4 | 6 | 10 |  |  |  |
| =================---=== | 5 | 5 | 10 |  |  |  |
| ============-------==== | 6 | 4 | 10 |  |  |  |
| =======-------------=== | 2 | 8 | 10 |  |  |  |
| ==================TTAGATCCAGT===== | 0 | 9 | 9 |  |  |  |
| ==================TATGCTTATGATTA===== | 0 | 9 | 9 |  |  |  |
| =====================-- | 0 | 8 | 8 |  |  |  |
| =====----------======== | 3 | 5 | 8 |  |  |  |
| ========--------------= | 0 | 8 | 8 |  |  |  |
| =================GC====== | 3 | 4 | 7 |  |  |  |
| =================CCGTCCA====== | 3 | 4 | 7 |  |  |  |
| ==========-----------== | 4 | 3 | 7 |  |  |  |
| ==================-==== | 2 | 4 | 6 |  |  |  |
| ====================--- | 2 | 4 | 6 |  |  |  |
| ================----=== | 3 | 3 | 6 |  |  |  |
| ========-------======== | 3 | 3 | 6 |  |  |  |
| ===================CGGGACAGTAC==== | 0 | 6 | 6 |  |  |  |
| ====================GTTAGGGATAGGGTTA=== | 3 | 3 | 6 |  |  |  |
| =======---------------= | 2 | 4 | 6 |  |  |  |
| =================T====== | 5 | 0 | 5 |  |  |  |
| =================A====== | 3 | 2 | 5 |  |  |  |
| ================-----== | 2 | 3 | 5 |  |  |  |
| =================------ | 1 | 4 | 5 |  |  |  |
| =============--------== | 0 | 5 | 5 |  |  |  |
| =======----------====== | 1 | 4 | 5 |  |  |  |
| ==========----------=== | 1 | 4 | 5 |  |  |  |
| =================-===== | 4 | 0 | 4 |  |  |  |
| =================TT====== | 1 | 3 | 4 |  |  |  |
| ================TC======= | 0 | 4 | 4 |  |  |  |
| ================--===== | 4 | 0 | 4 |  |  |  |
| ===========----======== | 3 | 1 | 4 |  |  |  |
| =================CGTCCA====== | 1 | 3 | 4 |  |  |  |
| ========---------====== | 1 | 3 | 4 |  |  |  |
| =============---------= | 1 | 3 | 4 |  |  |  |
| ========----------===== | 2 | 2 | 4 |  |  |  |
| =================GTGGGTTAGTAA====== | 0 | 4 | 4 |  |  |  |
| ===========------------ | 1 | 3 | 4 |  |  |  |
| =================ACACGACCTGGAA====== | 0 | 4 | 4 |  |  |  |
| ==========------------- | 1 | 3 | 4 |  |  |  |
| =================CGGGTTAGAGCCGTCCA====== | 1 | 3 | 4 |  |  |  |
| ================A======= | 3 | 0 | 3 |  |  |  |
| =================AC====== | 0 | 3 | 3 |  |  |  |
| ==============GG========= | 0 | 3 | 3 |  |  |  |
| =================GTT====== | 0 | 3 | 3 |  |  |  |
| ================GGA======= | 0 | 3 | 3 |  |  |  |
| ============CTC=====A====== | 0 | 3 | 3 |  |  |  |
| =============GCAC========== | 0 | 3 | 3 |  |  |  |
| =========------======== | 1 | 2 | 3 |  |  |  |
| =====-=======------==== | 0 | 3 | 3 |  |  |  |
| =================CAGTAACCCA====== | 0 | 3 | 3 |  |  |  |
| =========----------==== | 2 | 1 | 3 |  |  |  |
| ===============G======== | 0 | 2 | 2 |  |  |  |
| ==================C===== | 0 | 2 | 2 |  |  |  |
| ====================T=== | 0 | 2 | 2 |  |  |  |
| ================AA======= | 2 | 0 | 2 |  |  |  |
| ==============TA========= | 2 | 0 | 2 |  |  |  |
| ==================CT===== | 0 | 2 | 2 |  |  |  |
| =============--======== | 2 | 0 | 2 |  |  |  |
| ===================ACA==== | 2 | 0 | 2 |  |  |  |
| =================GGC====== | 0 | 2 | 2 |  |  |  |
| =================GTC====== | 0 | 2 | 2 |  |  |  |
| =================CGTC====== | 2 | 0 | 2 |  |  |  |
| =================CCCC====== | 0 | 2 | 2 |  |  |  |
| =================GACG====== | 0 | 2 | 2 |  |  |  |
| ====================GTCCA=== | 0 | 2 | 2 |  |  |  |
| =================CAAAAA====== | 1 | 1 | 2 |  |  |  |
| ======---=========--=== | 2 | 0 | 2 |  |  |  |
| =================ACGTCCA====== | 2 | 0 | 2 |  |  |  |
| ===============GTTCCCG======== | 0 | 2 | 2 |  |  |  |
| ==========--===----==== | 2 | 0 | 2 |  |  |  |
| ===============GAGCCAGT======== | 0 | 2 | 2 |  |  |  |
| ===============GG=======ATTCCA= | 0 | 2 | 2 |  |  |  |
| ===========TCTGAT===--======= | 0 | 2 | 2 |  |  |  |
| ===============CCGGGTCCCG======== | 0 | 2 | 2 |  |  |  |
| =================AACTGGGAAA====== | 0 | 2 | 2 |  |  |  |
| =----------============ | 2 | 0 | 2 |  |  |  |
| =================CCTGCCGTCCA====== | 0 | 2 | 2 |  |  |  |
| ====-----------======== | 1 | 1 | 2 |  |  |  |
| ======================ACAGGCGCGAAG= | 0 | 2 | 2 |  |  |  |
| =------------========== | 2 | 0 | 2 |  |  |  |
| =================CAGTTAGTCCAGTA====== | 2 | 0 | 2 |  |  |  |
| ========-------------== | 0 | 2 | 2 |  |  |  |
| =====--------------==== | 1 | 1 | 2 |  |  |  |
| ==============CACGTCCAGCACGTCC========= | 2 | 0 | 2 |  |  |  |
| =================AGTGTTTAACCAGTCTAACCA====== | 0 | 2 | 2 |  |  |  |
| ==================CTGGACTTAACTGGACTCTAAC===== | 2 | 0 | 2 |  |  |  |
| =====-============ACTCCTTAATTAGAGTTAAGT===== | 0 | 2 | 2 |  |  |  |
| =================TTGTAAAAAAAAGTAAAAAAAAA====== | 0 | 2 | 2 |  |  |  |
| ======================A= | 1 | 0 | 1 |  |  |  |
| ===================T==== | 1 | 0 | 1 |  |  |  |
| ==============C========= | 1 | 0 | 1 |  |  |  |
| ===============A======== | 0 | 1 | 1 |  |  |  |
| ===============C======== | 0 | 1 | 1 |  |  |  |
| ===============CG======== | 1 | 0 | 1 |  |  |  |
| ======================- | 1 | 0 | 1 |  |  |  |
| ===================TA==== | 1 | 0 | 1 |  |  |  |
| ================GA======= | 1 | 0 | 1 |  |  |  |
| ==================AT===== | 0 | 1 | 1 |  |  |  |
| ===================AG==== | 0 | 1 | 1 |  |  |  |
| ===================AT==== | 0 | 1 | 1 |  |  |  |
| ===================CA==== | 0 | 1 | 1 |  |  |  |
| ======================TA= | 0 | 1 | 1 |  |  |  |
| =====================TA== | 0 | 1 | 1 |  |  |  |
| ====================TA=== | 0 | 1 | 1 |  |  |  |
| =================GT====== | 0 | 1 | 1 |  |  |  |
| ===================TG==== | 0 | 1 | 1 |  |  |  |
| =================TA====== | 0 | 1 | 1 |  |  |  |
| ================GC======= | 0 | 1 | 1 |  |  |  |
| =========--============ | 1 | 0 | 1 |  |  |  |
| ============--========= | 1 | 0 | 1 |  |  |  |
| =================CAA====== | 1 | 0 | 1 |  |  |  |
| ===================CCA==== | 1 | 0 | 1 |  |  |  |
| ====================CTA=== | 1 | 0 | 1 |  |  |  |
| =====================CCA== | 1 | 0 | 1 |  |  |  |
| ===================TCA==== | 1 | 0 | 1 |  |  |  |
| ============GGG=========== | 0 | 1 | 1 |  |  |  |
| ============GGT=========== | 0 | 1 | 1 |  |  |  |
| ===============CCG======== | 0 | 1 | 1 |  |  |  |
| ===============CTC======== | 0 | 1 | 1 |  |  |  |
| ===============GGT======== | 0 | 1 | 1 |  |  |  |
| =================AAA====== | 0 | 1 | 1 |  |  |  |
| =================AAG====== | 0 | 1 | 1 |  |  |  |
| ==================ACC===== | 0 | 1 | 1 |  |  |  |
| ===================CCT==== | 0 | 1 | 1 |  |  |  |
| ====================CCA=== | 0 | 1 | 1 |  |  |  |
| ===================TCC==== | 0 | 1 | 1 |  |  |  |
| =================TAA====== | 0 | 1 | 1 |  |  |  |
| =================TTT====== | 0 | 1 | 1 |  |  |  |
| ================TCC======= | 0 | 1 | 1 |  |  |  |
| =============GGG========== | 0 | 1 | 1 |  |  |  |
| ===============CGGA======== | 1 | 0 | 1 |  |  |  |
| ==================GGGT===== | 1 | 0 | 1 |  |  |  |
| ===================GACG==== | 1 | 0 | 1 |  |  |  |
| =================TCCA====== | 1 | 0 | 1 |  |  |  |
| ===============GAGT======== | 0 | 1 | 1 |  |  |  |
| ================AACC======= | 0 | 1 | 1 |  |  |  |
| =================CCCA====== | 0 | 1 | 1 |  |  |  |
| =================CCCG====== | 0 | 1 | 1 |  |  |  |
| =================CGGT====== | 0 | 1 | 1 |  |  |  |
| =================GCCC====== | 0 | 1 | 1 |  |  |  |
| ===================TAGA==== | 0 | 1 | 1 |  |  |  |
| ===----================ | 1 | 0 | 1 |  |  |  |
| =================GTATT====== | 1 | 0 | 1 |  |  |  |
| ===================GAACC==== | 1 | 0 | 1 |  |  |  |
| =================TCCAA====== | 1 | 0 | 1 |  |  |  |
| =================CCCCA====== | 0 | 1 | 1 |  |  |  |
| =================CGCCC====== | 0 | 1 | 1 |  |  |  |
| =================CGTGT====== | 0 | 1 | 1 |  |  |  |
| =================GCCCA====== | 0 | 1 | 1 |  |  |  |
| =================GTCAA====== | 0 | 1 | 1 |  |  |  |
| ====================AGTTA=== | 0 | 1 | 1 |  |  |  |
| ===================TGTCA==== | 0 | 1 | 1 |  |  |  |
| =================CTGGAC====== | 1 | 0 | 1 |  |  |  |
| ==============TAGCT=======A== | 1 | 0 | 1 |  |  |  |
| =--===========--======= | 0 | 1 | 1 |  |  |  |
| =================AAGTAA====== | 0 | 1 | 1 |  |  |  |
| =================AGTCCA====== | 0 | 1 | 1 |  |  |  |
| =================CAGTAA====== | 0 | 1 | 1 |  |  |  |
| =================CCGTCC====== | 0 | 1 | 1 |  |  |  |
| ==================CAGTAA===== | 0 | 1 | 1 |  |  |  |
| =================GGGCAA====== | 0 | 1 | 1 |  |  |  |
| ===================AATAAT==== | 0 | 1 | 1 |  |  |  |
| ===================ACTACA==== | 0 | 1 | 1 |  |  |  |
| ===================TAATTA==== | 0 | 1 | 1 |  |  |  |
| ===================TAGTTA==== | 0 | 1 | 1 |  |  |  |
| =================TAACCA====== | 0 | 1 | 1 |  |  |  |
| ================GTGTTA======= | 0 | 1 | 1 |  |  |  |
| ==============GA=======AGTA== | 0 | 1 | 1 |  |  |  |
| =============TGCG====TG====== | 0 | 1 | 1 |  |  |  |
| ===========-====----=== | 1 | 0 | 1 |  |  |  |
| =================TGGACCA====== | 1 | 0 | 1 |  |  |  |
| =================AGTCCAA====== | 0 | 1 | 1 |  |  |  |
| =================CCAGTAT====== | 0 | 1 | 1 |  |  |  |
| =================TAACCCA====== | 0 | 1 | 1 |  |  |  |
| =================TGGGAAA====== | 0 | 1 | 1 |  |  |  |
| ================TGGAGTA======= | 0 | 1 | 1 |  |  |  |
| =============GGGAACT========== | 0 | 1 | 1 |  |  |  |
| =============TAGACGT========== | 0 | 1 | 1 |  |  |  |
| ==------=========C====== | 1 | 0 | 1 |  |  |  |
| ======----========--=== | 1 | 0 | 1 |  |  |  |
| =================AGTAATAA====== | 1 | 0 | 1 |  |  |  |
| ==============GAGTTAGT========= | 1 | 0 | 1 |  |  |  |
| =============T====CTGGCCT====== | 1 | 0 | 1 |  |  |  |
| ======C=======------==== | 0 | 1 | 1 |  |  |  |
| =======-------========= | 0 | 1 | 1 |  |  |  |
| =================CAGTCCAA====== | 0 | 1 | 1 |  |  |  |
| ==================GCCTGGCT===== | 0 | 1 | 1 |  |  |  |
| ==-==========------==== | 1 | 0 | 1 |  |  |  |
| =========-----====--=== | 1 | 0 | 1 |  |  |  |
| ===T=======-------====== | 1 | 0 | 1 |  |  |  |
| ==-=======------======= | 0 | 1 | 1 |  |  |  |
| ================AAAGCTCCG======= | 0 | 1 | 1 |  |  |  |
| =================AGTAATGAA====== | 0 | 1 | 1 |  |  |  |
| =================CAGTAACCA====== | 0 | 1 | 1 |  |  |  |
| =================CAGTCCAGT====== | 0 | 1 | 1 |  |  |  |
| =================CCCGTCCAA====== | 0 | 1 | 1 |  |  |  |
| ==================ATAACCAGT===== | 0 | 1 | 1 |  |  |  |
| ===================AATAATAAT==== | 0 | 1 | 1 |  |  |  |
| ======---------======== | 1 | 0 | 1 |  |  |  |
| =======GT====-------===== | 1 | 0 | 1 |  |  |  |
| =============GTTTCGTTCC========== | 1 | 0 | 1 |  |  |  |
| ==================AGTAATAAGC===== | 0 | 1 | 1 |  |  |  |
| ==================GGTCCAGTTA===== | 0 | 1 | 1 |  |  |  |
| ====================GTTACTACTA=== | 0 | 1 | 1 |  |  |  |
| =================TAGATTACTA====== | 0 | 1 | 1 |  |  |  |
| =================TGTGGTTCCA====== | 0 | 1 | 1 |  |  |  |
| =================GACGGTTAGTA====== | 0 | 1 | 1 |  |  |  |
| ==============CGGGACGGG=====AC==== | 0 | 1 | 1 |  |  |  |
| ============ACGGGACGTTAG=========== | 1 | 0 | 1 |  |  |  |
| =================CCACGCCAGTAC====== | 1 | 0 | 1 |  |  |  |
| =================CGTCCAGTAAAA====== | 1 | 0 | 1 |  |  |  |
| =================GTGTTAGTGTTA====== | 0 | 1 | 1 |  |  |  |
| =================TATATAAGTCCA====== | 0 | 1 | 1 |  |  |  |
| ----====-------======== | 1 | 0 | 1 |  |  |  |
| ==-==========---------- | 1 | 0 | 1 |  |  |  |
| ======------------===== | 1 | 0 | 1 |  |  |  |
| =----=====-------====== | 0 | 1 | 1 |  |  |  |
| ======----===-------=== | 0 | 1 | 1 |  |  |  |
| =======------------==== | 0 | 1 | 1 |  |  |  |
| ========------------=== | 0 | 1 | 1 |  |  |  |
| =================ACACTAATCGTGT====== | 0 | 1 | 1 |  |  |  |
| =================CAGAGATCGCTGC====== | 0 | 1 | 1 |  |  |  |
| ================TAGAGCTAGAAGT======= | 0 | 1 | 1 |  |  |  |
| =----------=======--=== | 1 | 0 | 1 |  |  |  |
| =====-------------===== | 0 | 1 | 1 |  |  |  |
| =========-------------= | 0 | 1 | 1 |  |  |  |
| ===============GTTAGTTAGTTAGT======== | 0 | 1 | 1 |  |  |  |
| =================CCATGATTCTAACC====== | 0 | 1 | 1 |  |  |  |
| =================ACCGGTTCCCGTCCA====== | 1 | 0 | 1 |  |  |  |
| ====================TTCCCAGTCTACCCC=== | 1 | 0 | 1 |  |  |  |
| ====--------------===== | 0 | 1 | 1 |  |  |  |
| =================AAGTAGTAAAGTAAA====== | 0 | 1 | 1 |  |  |  |
| =================ATAACCAGTAACCAA====== | 0 | 1 | 1 |  |  |  |
| =================GGCTGTGGTATATAT====== | 0 | 1 | 1 |  |  |  |
| =================GTGGGACAGAGAACA====== | 0 | 1 | 1 |  |  |  |
| --------=====------==== | 1 | 0 | 1 |  |  |  |
| ====================GGTAGGGATAGGGTTA=== | 1 | 0 | 1 |  |  |  |
| ===============-------AAAAAAAA= | 0 | 1 | 1 |  |  |  |
| ====================GTTAGGGCTAGGGTTA=== | 0 | 1 | 1 |  |  |  |
| ==========TTTCCACA=====-------= | 0 | 1 | 1 |  |  |  |
| =================GTCTGGTATAGTCCAGA====== | 1 | 0 | 1 |  |  |  |
| ==========-----==GTGCGAATCTG====== | 0 | 1 | 1 |  |  |  |
| =================CCCCTCGGGGTTGGGAG====== | 0 | 1 | 1 |  |  |  |
| ==================GAATTAGATAAA====AGTTT= | 0 | 1 | 1 |  |  |  |
| ====-----------------== | 1 | 0 | 1 |  |  |  |
| ============-====ACACGACCTGGATGGA====== | 1 | 0 | 1 |  |  |  |
| =================CAGTACTGGCTGCAGTAC====== | 0 | 1 | 1 |  |  |  |
| =================CCGCCGCGTGCGAATCTG====== | 0 | 1 | 1 |  |  |  |
| ==================GATTAGTAATTAGAGTAA===== | 0 | 1 | 1 |  |  |  |
| ==================GTAATTAGTAATTAGTAA===== | 0 | 1 | 1 |  |  |  |
| =================AAGTAAGTTAAGTAAGTAA====== | 1 | 0 | 1 |  |  |  |
| ==============GATATATATATATATATAT========= | 0 | 1 | 1 |  |  |  |
| =================CCGTCCAGTAAAAAAAGTAA====== | 0 | 1 | 1 |  |  |  |
| ===================AAGACTATCTTGCCAAAAAA==== | 0 | 1 | 1 |  |  |  |
| ============GTTAGAGCTAGAAA======ACCTGGA===== | 1 | 0 | 1 |  |  |  |
| =================CAAAAGCTCGAGGTGGGGGAG====== | 0 | 1 | 1 |  |  |  |
| =================GGTAGACGGTAGTCCAGTAGA====== | 0 | 1 | 1 |  |  |  |
| =================TTGTAAAAAAAAGTAAAAAAA====== | 0 | 1 | 1 |  |  |  |
| ======CGTCCAGTAATGGTTATGTTAT================= | 0 | 1 | 1 |  |  |  |
| =================AAGTAAGTAAGTTAAGTAAGTAA====== | 1 | 0 | 1 |  |  |  |
| ==================GACGTTACTAATTAGGGTTAATT===== | 0 | 1 | 1 |  |  |  |
| ================TAAGTAAGGTGGCGCGGGGTAAA======= | 0 | 1 | 1 |  |  |  |
| ===================TTGGGTTAGGGTTAGGGTTAGGGTTA==== | 1 | 0 | 1 |  |  |  |
| =================TGGACGGGGTAAACTGGGAAAGTGAT====== | 0 | 1 | 1 |  |  |  |
| ================TAAGTAAGTAAGTAAGTAAGTAAGTA======= | 0 | 1 | 1 |  |  |  |
